# Antibacterial nucleoside-analog inhibitor of bacterial RNA polymerase: pseudouridimycin

**DOI:** 10.1101/106906

**Authors:** Sonia I. Maffioli, Yu Zhang, David Degen, Thomas Carzaniga, Giancarlo Del Gatto, Stefania Serina, Paolo Monciardini, Carlo Mazzetti, Paola Guglierame, Gianpaolo Candiani, Alina Iulia Chiriac, Giuseppe Facchetti, Petra Kaltofen, Hans-Georg Sahl, Gianni Dehò, Stefano Donadio, Richard H. Ebright

## Abstract

There is an urgent need for new antibacterial drugs effective against bacterial pathogens resistant to current drugs^1–2^. Nucleoside-analog inhibitors (NAIs) of viral nucleotide polymerases have had transformative impact in treatment of HIV^3^ and HCV^4^. NAIs of bacterial RNA polymerase (RNAP) potentially could have major impact on treatment of bacterial infection, particularly because functional constraints on substitution of RNAP nucleoside triphosphate (NTP) binding sites^4-5^ could limit resistance emergence^4-5^. Here we report the discovery, from microbial extract screening, of an NAI that inhibits bacterial RNAP and exhibits antibacterial activity against a broad spectrum of drug-sensitive and drug-resistant bacterial pathogens: pseudouridimycin (PUM). PUM is a novel microbial natural product consisting of a formamidinylated, N-hydroxylated Gly-Gln dipeptide conjugated to 6'-amino-pseudouridine. PUM potently and selectively inhibits bacterial RNAP in vitro, potently and selectively inhibits bacterial growth in culture, and potently clears infection in a mouse model of *Streptococcus pyogenes* peritonitis. PUM inhibits RNAP through a binding site on RNAP (the "i+1" NTP binding site) and mechanism (competition with UTP for occupancy of the "i+1" NTP binding site) that differ from those of the RNAP inhibitor and current antibacterial drug rifampin (Rif). PUM exhibits additive antibacterial activity when co-administered with Rif, exhibits no cross-resistance with Rif, and exhibits a spontaneous resistance rate an order-of-magnitude lower than that of Rif. The results provide the first example of a selective NAI of bacterial RNAP, provide an advanced lead compound for antibacterial drug development, and provide structural information and synthetic routes that enable lead optimization for antibacterial drug development.

---

We screened a library of 3000 Actinobacterial^6^ and fungal culture extracts for inhibition of RNAP, and we identified two extracts that inhibited bacterial RNAP (*E. coli* RNAP) but did not inhibit a structurally unrelated bacteriophage RNAP (SP6 RNAP) and did not contain a previously characterized inhibitor of bacterial RNAP (see Methods). Fractionation of the two extracts by reversed-phase chromatography and structure elucidation of active components by mass spectrometry and multidimensional NMR spectrometry revealed that the extracts contained the same novel active component: pseudouridimycin (PUM; Fig. 1a; Extended Data Figs. 1-2).

**Figure 1.**
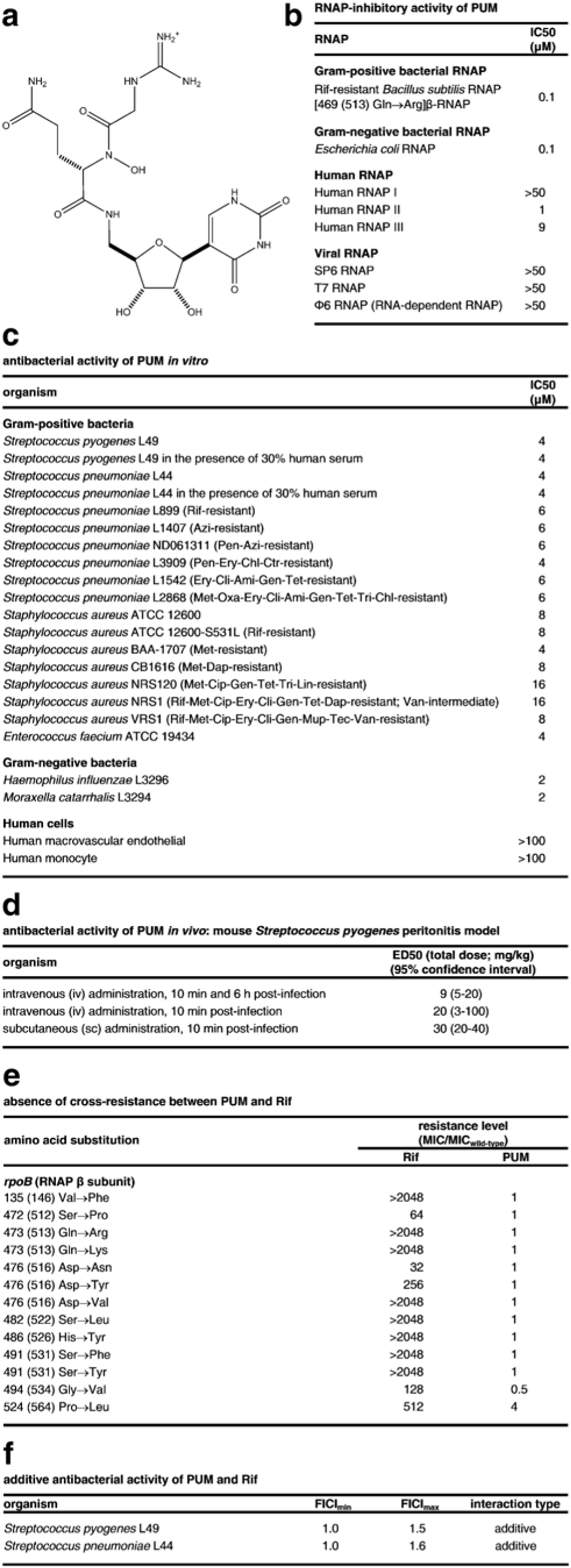
**Structure, RNAP-inhibitory activity, and antibacterial activity of PUM. a**, Structure of PUM. **b**, RNAP-inhibitory activity of PUM. **c**, Antibacterial activity of PUM *in vitro*. **d**, Antibacterial activity of PUM *in vivo*. Drug resistances are as follows: Ami. amikacin; Azi, azithromycin; Cip, ciprofloxacin; Ctr, ceftriaxone; Dap, daptomycin; Ery, erythromycin; Chl, chloramphenicol; Cli, clindamycin; Gen, gentamycin; Lin, linezolid; Met, methicillin; Mup, mupirocin; Pen, penicillin; Ox, oxacillin; Rif, rifampin; Tec, teicoplanin; Tet, tetracycline; Tri, trimethoprim; Van, vancomycin. **e**, Absence of cross-resistance between PUM and Rif (data for *S. pyogenes* Rif-resistant mutants; residues numbered as in *S. pyogenes* and, in parentheses, *E. coli*). **f**, Additive antibacterial activity of PUM and Rif.

PUM selectively inhibits bacterial RNAP (IC50 = 0.1 μM; selectivity >4- to >500-fold; Fig. 1b; Extended Data Table 1; Extended Data Fig. 3), selectively inhibits bacterial growth (IC50 = 2 to 16 μM; selectivity >6- to >60-fold; Fig. 1c), and clears infection *in vivo* in a mouse *Streptococcus pyogenes* peritonitis model (ED50 = 9 mg/kg; Fig. 1c; Extended Data Table 2). PUM exhibits antibacterial activity against both Gram-positive and Gram-negative bacteria and against both drug-sensitive and drug-resistant bacterial strains, including rifamycin-, β-lactam-, fluoroquinolone- macrolide-, tetracycline-, aminoglycoside-, lincosamide-, chloramphenicol-, oxazolidinone-, trimethoprim-, glycopeptide-, lipopeptide-, mupirocin-, and multi-drug-resistant strains (Fig. 1c).

PUM exhibits no cross-resistance with the classic RNAP inhibitor Rif (Figs. 1b-c,e), exhibits additive antibacterial activity when co-administered with Rif (Fig. 1f), and exhibits spontaneous resistance rates an order-of-magnitude lower than those of Rif (Fig. 2a), suggesting that PUM inhibits RNAP through a binding site and mechanism different from those of Rif.

**Figure 2.**
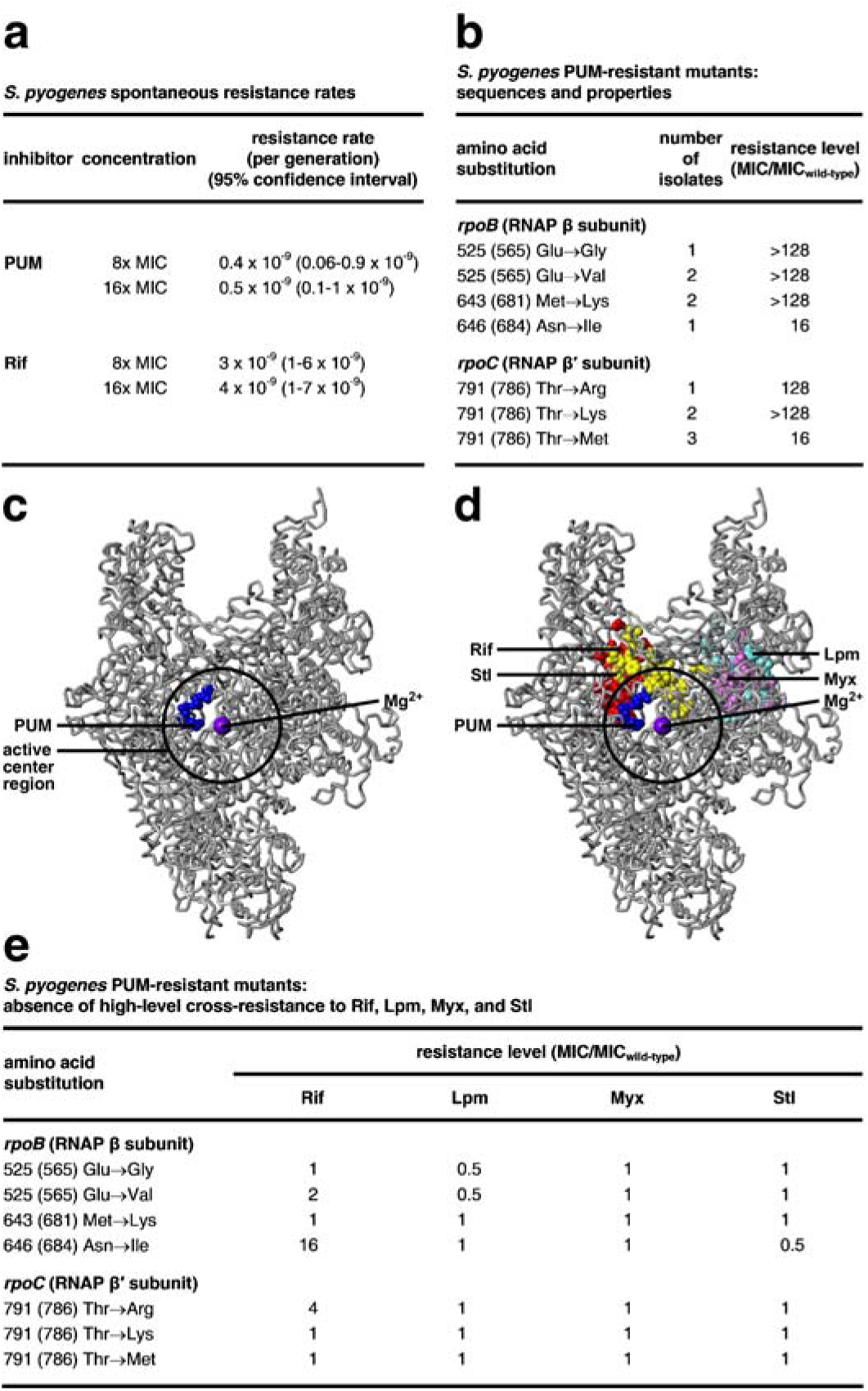
**Target of PUM: RNAP i+1 NTP binding site. a**, Spontaneous resistance rates for PUM and Rif. **b**, *S. pyogenes*spontaneous PUM-resistant mutants. **c**, Location of PUM target (blue) in three-dimensional structure of bacterial RNAP^12^ (gray; black circle for active-center region; violet sphere for active-center Mg^2+^(I); β′ non-conserved region and σ omitted for clarity). **d**, Absence of overlap between PUM target (blue) and Rif (red), Lpm (cyan), Myx (pink), and Stl (yellow) targets. **e**, Absence of high-level cross-resistance for *S. pyogenes* PUM-resistant mutants to Rif, Lpm, Myx, and Stl.

Gene sequencing indicates that PUM-resistant mutants contain mutations in the *rpoB* gene (encodes RNAP β subunit) or the *rpoC* gene (encodes RNAP β′ subunit), confirming that RNAP is the functional cellular target of PUM (Fig. 2b; Extended Data Fig. 4a-b). In the Gram-positive bacterium *S. pyogenes*, substitutions conferring ≥4x PUM-resistance are obtained at four sites: β residues 565, 681, and 684, and β′ residue 786 (numbered as in *E. coli* RNAP; Fig. 2b). In the Gram-negative bacterium *E. coli*, substitutions conferring PUM-resistance are obtained at two sites: β residues 565 and 681 (Extended Data Fig. 4a-b). The number of sites of substitutions conferring PUM-resistance is an order-of-magnitude lower than the number of sites of substitutions conferring Rif-resistance (2 to 4 vs. 25^7-8^), consistent with, and accounting for, the observation that spontaneous resistance rates for PUM are an order-of-magnitude lower than those for Rif (Fig. 2a).

Mapping the sites of substitutions conferring PUM resistance onto the three-dimensional structure of bacterial RNAP shows that the sites form a single discrete cluster ("PUM target"; Fig. 2c; Extended Data Fig. 4d). The PUM target is located within the RNAP active-center region and overlaps the RNAP active-center i+1 NTP binding site (Fig. 2c; Extended Data Fig. 4d), suggesting that PUM inhibits RNAP by interfering with function of the i+1 site. The PUM target is different from, and does not overlap, the Rif target (Fig. 2d; Extended Data Fig. 4e)^7-9^, consistent with, and accounting for, the observation that PUM does not share cross-resistance with Rif (Figs. 1b-c,e, 2e; Extended Data Fig. 4f-g) and the observation that PUM and Rif exhibit additive antibacterial activity (Fig. 1f). The PUM target also is different from, and does not overlap, the targets of the RNAP inhibitors lipiarmycin (Lpm)^10-11^, myxopyronin (Myx)^11-13^, streptolydigin (Stl)^14-15^, CBR703 (CBR)^16-17^, and salinamide (Sal)^18^ (Fig. 2d; Extended Data Fig. 4e), and, correspondingly, PUM does not exhibit cross-resistance with Lpm, Myx, Stl, CBR703, and Sal (Fig. 2e; Extended Data Fig. 4f,h-l). The PUM target partly overlaps the target for the RNAP inhibitor GE23077 (GE; Extended Data Fig. 4m)^5^, and, correspondingly, PUM exhibits partial cross-resistance with GE (Extended Data Fig. 4n).

The observation that PUM is an NAI that has the same Watson-Crick base-pairing specificity as UTP (Fig. 1a) and the observation that the PUM target overlaps the RNAP i+1 NTP binding site (Fig. 2b; Extended Data Fig. 4a) suggest the hypothesis that PUM functions as an NAI that competes with UTP for occupancy of the RNAP i+1 NTP binding site. Five biochemical results support this hypothesis. First, PUM inhibits transcription by inhibiting nucleotide addition (Extended Data Fig. 5). Second, high concentrations of UTP––but not high concentrations of GTP, ATP, or CTP––overcome transcription inhibition by PUM (Fig. 3a). Third, PUM inhibits transcription only on templates that direct incorporation of U (Fig. 3b). Fourth, in single-nucleotide-addition transcription reactions, PUM inhibits incorporation of U, but not G, A, or C (Fig. 3c). Fifth, in multiple-nucleotide-addition transcription reactions, PUM inhibits incorporation of U, but not G, A, or C (Fig. 3d). The results in Fig. 3c-d further establish that transcription inhibition by PUM not only requires a template position that directs incorporation of U, but also strongly prefers a preceding template position that directs incorporation of G, A, or U. We conclude that PUM functions as an NAI that competes with UTP at positions that direct incorporation of U preceded by positions that direct incorporation of G, A, or U.

**Figure 3.**
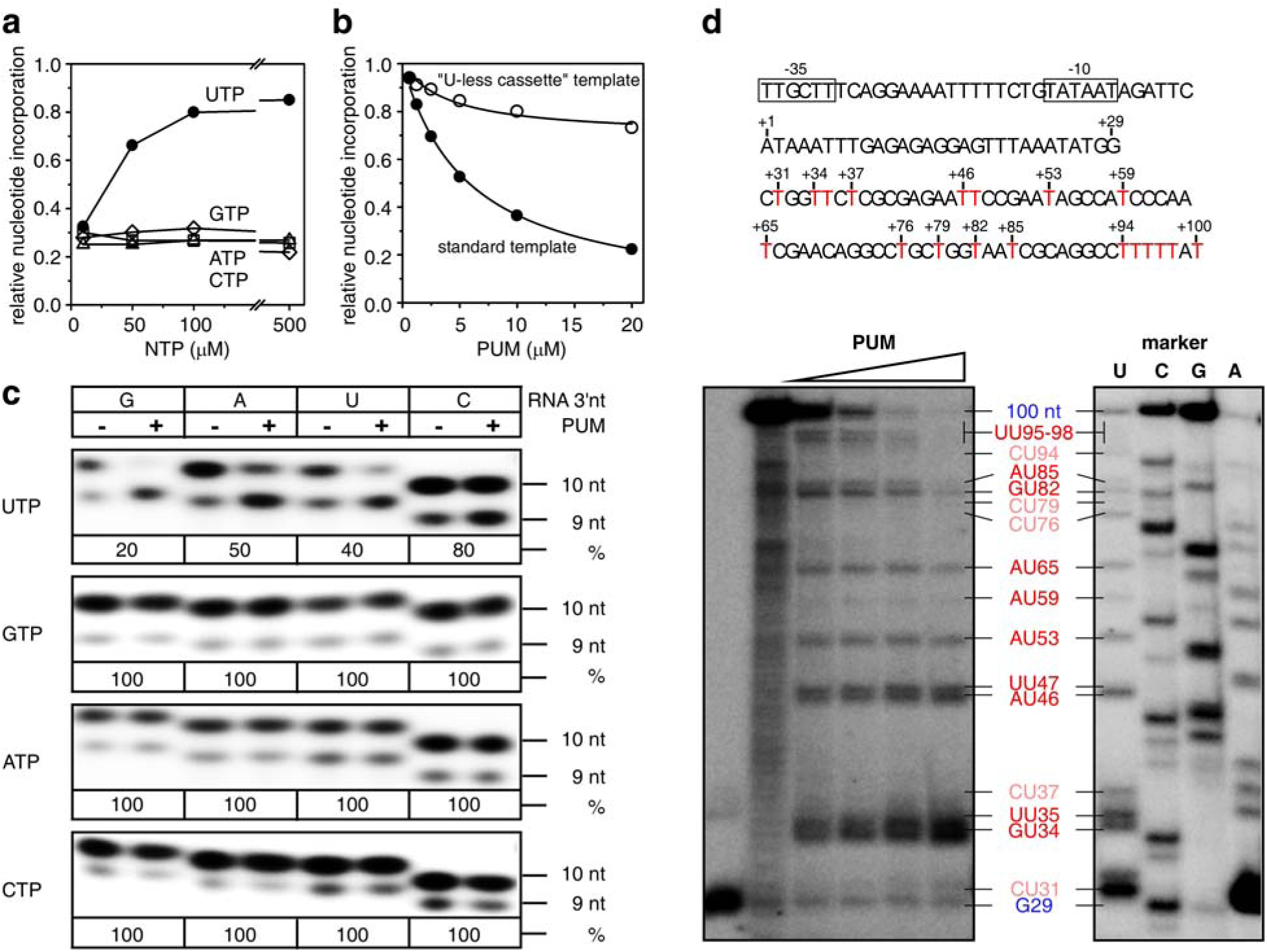
**Mechanism of PUM: competition with UTP for occupancy of RNAP i+1 NTP binding site. a**, Suppression of inhibition by PUM by high [UTP], but not high [GTP], [ATP], or [CTP]. **b**, Inhibition by PUM of transcription directing incorporation of U+G+A+C, but not "U-less" transcription specifying incorporation of G+A+C. **c**, Single-nucleotide-addition reactions showing that inhibition by PUM requires template positions directing incorporation of U (row 1) and prefers preceding template positions directing incorporation of G, A, or U (columns 1-3). **d**, Multiple-nucleotide-addition reactions showing that inhibition by PUM requires template positions directing incorporation of U (red GU, AU, or UU, and pink CU) and prefers preceding template positions directing incorporation of G, A, or U (red GU, AU, or UU).

To define the structural basis of transcription inhibition by PUM, we determined a crystal structure of a transcription initiation complex containing PUM (RPo-GpA-PUM; Fig. 4a) and, for comparison, a crystal structure of a corresponding transcription initiation complex containing CMPcPP (RPo-GpA-CMPcPP; Fig. 4b). The results establish that PUM is an NAI that competes for occupancy of the i+1 site (Fig. 4). PUM binds to the i+1 site (Fig. 4a). The PUM base makes Watson-Crick H-bonds with a DNA template-strand A in a manner equivalent to an NTP base; the PUM sugar moiety makes interactions with the i+1 site in a manner nearly equivalent to an NTP sugar; the PUM glutamine moiety makes interactions that mimic interactions made by an NTP triphosphate; and the PUM N-hydroxy and guanidinyl moieties interact with the RNA nucleotide base-paired to the preceding template position (RNA 3'-nucleotide), with the N-hydroxy donating an H-bond to the 3'-OH of the RNA 3'-nucleotide, and the guanidinyl moiety donating one H-bond to the 5'-phosphate of the RNA 3'-nucleotide and another to the base of the RNA 3'-nucleotide (Fig. 4a).

**Figure 4.**
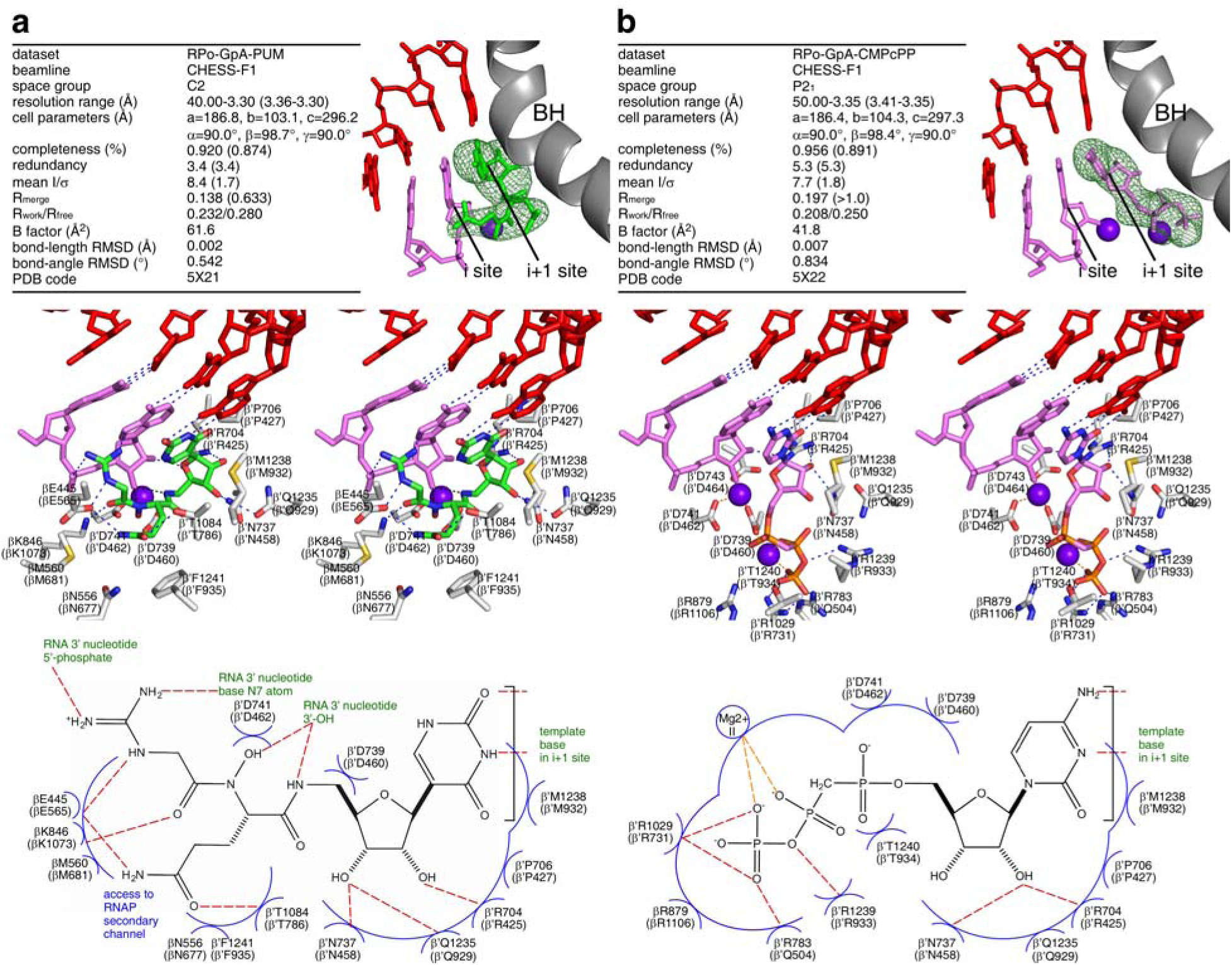
**Structural basis of transcription inhibition by PUM** Structures of transcription initiation complexes containing PUM (**a**) and CMPcPP (**b**). Top, crystallization and refinement statistics (left) and experimental electron density and fit (right). Green, PUM; pink, RNA and CMPcPP; red, DNA template strand; violet sphere between i and i+1 sites, Mg^2+^(I); violet sphere in i+1 site, Mg^2+^(II); gray; RNAP bridge helix; green mesh, mF_o_-DF_c_ omit map (contoured at 2.5σ). Middle, stereodiagram of interactions. Green, PUM carbon atoms; pink, RNA and CMPcPP carbon atoms; gray, RNAP carbon atoms; red, blue, yellow, and orange, oxygen, nitrogen, sulfur, and phosphorous atoms; dashed lines, H-bonds; other colors, as above. Bottom, summary of interactions. Red dashed lines, H-bonds; blue arcs, van der Waals interactions. Residues numbered as in *T. thermophilus* RNAP and, in parentheses, *E. coli* RNAP.

The structure of the PUM-inhibited complex accounts for the observed specificity of inhibition for template positions that direct incorporation of U preceded by template positions that direct incorporation of G, A, or U. The Watson-Crick base pair by the PUM base moiety provides absolute specificity for a position directing incorporation of U (Fig. 4a). The H-bond donated by the PUM guanidinyl moiety with the base of the RNA 3'-nucleotide confers specificity for a preceding position directing incorporation of G, A, or U (each of which contains an H-bond acceptor at the appropriate position; Fig. 4a).

The structure also explains the selectivity of transcription inhibition by PUM. All RNAP residues contacted by PUM are highly conserved across Gram-positive and Gram-negative bacterial RNAP (Extended Data Fig. 6), accounting for the inhibition of both Gram-positive and Gram-negative bacterial RNAP. In contrast, four RNAP residues important for PUM are not conserved in human RNAP I, II, and III (β residues 677, 681, and 684, and β′ residue 932; Extended Data Fig. 6), accounting for selectivity for bacterial RNAP over human RNAP I, II, and III.

The structure also explains the small size of the PUM resistance spectrum (four residues in *S. pyogenes* RNAP; two residues in *E. coli* RNAP; Fig. 2b; Extended Data Fig. 4a-b). PUM makes direct contacts with RNAP residues at which PUM-resistant substitutions are obtained (Extended Data Fig. 6). However, PUM also makes direct contacts with ten other RNAP residues that comprise functionally critical residues of the RNAP active center that cannot be substituted without compromising RNAP activity^5;19-23^, and thus which cannot be substituted to yield viable resistant mutants^5^ (Fig. 4a). We infer that PUM interacts with a *"privileged target"* for which most residues (10-12 of 14 residues) have functional constraints that limit substitution to yield viable resistant mutants. Similar results have been reported for the RNAP inhibitor GE, a non-nucleoside-analog inhibitor that binds to the RNAP i and i+1 sites^5^ and that exhibits a small target-based resistance spectrum^5^ (but that, unlike PUM, exhibits high non-target-based resistance, presumably at the level of uptake or efflux, precluding development as an antibacterial drug).

The structure enables structure-based design of novel PUM analogs with increased potency and increased selectivity. Initial lead-optimization efforts corroborate the importance of the PUM N-hydroxy, glutamine, and guanidinyl moieties and demonstrate that the PUM glutamine C(O)NH_2_ can be replaced by C(O)NHR while retaining RNAP inhibitory and antibacterial activity (Extended Data Fig. 7).

Our results provide a new class of antibiotic with activity against Gram-positive and Gram-negative bacteria *in vitro* and *in vivo*, no cross-resistance with current antibacterial drugs, and low rates of resistance emergence. Our discovery of a new class of antibiotic from conventional microbial extract screening indicates that, contrary to widespread belief^1^, conventional microbial extract screening has not been exhausted as a source of new antibacterial lead compounds.

Our results provide the first example of a selective NAI of bacterial RNAP. NAIs of viral nucleotide polymerases have been of immense importance for development of anti-HIV^3^ and anti-HCV^4^ drugs. NAIs of bacterial RNAP may show comparable promise for development of antibacterial drugs.

## Acknowledgements

Work was supported by NIH grants GM041376 and AI104660 to RHE and Italian Ministry of Research and Regione Lombardia grants to NAICONS. We thank former colleagues at Vicuron Pharmaceuticals for early characterization of PUM, S. Parapini for cytotoxicity assays, W. Fenical for Sal, A. Berk and I. Grummt for plasmids, NARSA and BEI Resources for strains, and the Cornell High Energy Synchrotron Source for beamline access.

## Data Availability

Atomic coordinates and structure factors for reported crystal structures have been deposited with the Protein Data Bank under accession codes 5X21 (RPo-GpA-PUM) and 5X22 (RPo-GpA-CMPcPP).

## Author Contributions

S.M. performed structure elucidation of PUM and semi-syntheses and syntheses of PUM derivatives. Y.Z. purified RNAP, assessed template-specificity of inhibition and determined crystal structures. D.D. purified RNAP, and assessed RNAP-inhibitory activities, antibacterial activities, resistance, and cross-resistance. T.C., S.S., and G.D. assessed RNAP-inhibitory activities. G.D.G. and C.M. purified PUM and participated in semi-syntheses and syntheses of PUM derivatives. P.M. performed characterization and fermentation of the PUM producer strain. P.G. and G.C. performed mouse infection studies. A.I.C. and H.-G.S. performed macromolecular synthesis assays. G.F. and P.K. performed microbial extract screening and dereplication. S.D. and R.H.E. designed the study, analyzed data, and wrote the paper.

## Author Information

S.M., S.S, P.M., and S.D. are employees and/or shareholders of NAICONS. Y.Z., S.M., S.S., P.M., P.G., G.C., S.D., and R.H.E. have patent filings. The other authors declare that no competing interests exist. (Vicuron Pharmaceuticals no longer is operational.) Correspondence and requests for materials should be addressed to S.D. or R.H.E.

## METHODS

### *E. coli* RNAP core enzyme

For experiments in Fig. 3c, *E. coli* RNAP core enzyme was prepared from *E. coli* strain BL21(DE3) (Invitrogen) transformed with plasmids pEcABC-H6^24^ and pCDFω^25^, using procedures as described^26^

For experiments assessing promoter-independent transcription in Extended Data Table 1, *E. coli* RNAP core enzyme was prepared from *E. coli* strain XE54^27^ transformed with plasmid pRL706^28^, using procedures as described^29^.

### *E.coli* RNAP σ^70^ holoenzyme

For experiments in Fig. 3d, *E. coli* RNAP σ^70^ holoenzyme was prepared from *E. coli* strain XE54^26^ transformed with plasmid pREII-NHα^29^, using procedures as described^18^.

For experiments in Extended Data Fig. 4c, *E. coli* RNAP σ^70^ holoenzyme and [565Glu→Asp]β-RNAP σ^70^ holoenzyme were prepared from *E. coli* strain XE54^26^ transformed with plasmid pRL706^28^ or pRL706-565D^5^, using procedures as described^29^.

### *B.subtilis* RNAP σ^A^ holoenzyme

Rif-resistant *B. subtilis* [469 (513) Gln→Arg]β-RNAP σ^A^ holoenzyme was prepared from *B. subtilis* strain MH5636-Q469R (spontaneous Rif-resistant mutant of *B. subtilis* strain MH5636^30^; selected on LB agar containing 2 μg/ml Rif; confirmed by PCR amplification and sequencing of *rpoB*) using procedures essentially as described^30^.

### T. thermophilus RNAP σ^A^ holoenzyme

*T. thermophilus* RNAP σ^A^ holoenzyme was prepared as described^5^.

### Microbial extract screening

A sub-library of ~3,000 microbial extracts (prepared as described^31^) with growth-inhibitory activity against *Staphylococcus aureus* ATCC 6538 was screened for the ability to inhibit *E. coli* RNAP and bacteriophage SP6 RNAP (Promega). Screening was performed using 96-well microplates. Reactions contained (50 µ1): 5 µ1 extract (dissolved in 10% DMSO), 0.2 U *E. coli* RNAP σ^70^ holoenzyme (Sigma-Aldrich) or 0.2 U SP6 RNAP (Promega), 0.2 nM plasmid pUC18 (Clontech; for assays with *E. coli* RNAP) or 0.2 nM plasmid pGEM-3Z (Promega; for assays with SP6 RNAP), 500 µM ATP, 500 µM GTP, 500 µM CTP, and 2 µM [α^32^P]UTP (0.2 Bq/fmol; PerkinElmer), in 20 mM Tris-acetate (pH 7.9), 50 mM KCl, 4 mM magnesium acetate, 0.1 mM EDTA, 5 mM dithiothreitol, and 100 µg/ml bovine serum albumin. Reaction components except DNA were pre-equilibrated 10 min at 22°C. Reactions were initiated by addition of DNA, were allowed to proceed 1 h at 22°C, and were terminated by addition of 150 µl ice-cold 10% (w/v) trichloroacetic acid (TCA). After 1 h at 4°C, resulting TCA precipitates were collected on glass-fiber filters (UniFilter GF/B; PerkinElmer) using a 96-well harvester (Packard) and were washed once with water. Radioactivity was quantified using a TopCount scintillation counter (Packard), and % inhibition was calculated as:

% inhibition = 100-[100(R_sample_-R_neg_)/(R_pos_-R_neg_)]

where R_sample_, R_pos_ and R_neg_ are observed radioactivity levels in a reaction, in a control reaction without extract, and in a control reaction without plasmid, respectively.

Two extracts that inhibited the reaction with *E. coli* RNAP by ≥80%, that did not inhibit the reaction with SP6 RNAP, and that did not contain mass-spectrometry signals indicative of a previously characterized RNAP inhibitor, were designated as "hit extracts."

### Characterization of producer strains

The producer strains of the hit extracts were strains IDI38640 and IDI38673. ID38640 and ID38673 are Actinobacterial isolates from soil samples collected in Italy and France, respectively. ID38640 and ID38673 exhibit cell morphologies consistent with the genus *Streptomyces* and exhibit 16S rRNA gene sequences (determined as described ^32^; GenBank accession numbers JQ929050 and JQ929051) that were 99.9% identical over 1.4 kB to each other and were highly similar to those of a cluster of closely-related *Streptomyces* species (*S. nigrescens*, *S. tubercidicus*, *S. rimosus* subsp. *rimosus*, *S. hygroscopicus* subsp. *angustmyceticus*, and *S. libani* subsp. *libani*).

### Isolation and purification of PUM

For each producer strain of a hit extract, the strain was cultured on a 55 mm BTT agar plate^31^ 4-7 days at 30°C, the mycelium was scraped from the plate and used to inoculate a 50 ml Erlenmeyer flask containing 15 ml of seed medium (20 g/L dextrose monohydrate, 2 g/L yeast extract, 8 g/L soybean meal, 1 g/L NaCl, and 4 g/L CaCO_3_, pH 7.3), and the resulting culture was incubated 48 h at 30°C on a rotary shaker (200 rpm agitation) for 48 h. Following initial incubation, 5 ml of the culture was used to inoculate 100 ml of fresh seed medium in a 500-ml flask, and the resulting culture was incubated 72 h under the same conditions. A 5% (v/v) inoculum was transferred into 2 L of production medium (10 g/L dextrose monohydrate, 24 g/L maize dextrin, 8 g/L soy peptone, 5 g/L yeast extract, and 1 g/L NaCl, pH 7.2) in a 3-L vessel, and the resulting culture was grown in a BioFlo 115 Fermentor (Eppendorf) 96 h at 30°C, with aeration at 0.5 volume air per volume medium per min stirring at 600 rpm. The culture was filtered through 10.25 in disc filter paper (Scienceware), and the resulting cleared broth was concentrated to ~1 L *in vacuo* and loaded onto a column of 500 mg of Dowex 50W x 400 mesh (previously activated with two bed volumes of 5% HCl and washed with H_2_O until neutralization). After washing with 5 bed volumes each of 20 mM sodium acetate at pH 6 and sodium acetate at pH 7, PUM was eluted using six bed volumes of 100 mM NH_4_OAc at pH 9. PUM-containing fractions were desalted by reversed-phase medium-pressure liquid chromatography on Combiflash Rf (Teledyne Isco) using a 30 g C18 RediSep Rf column (Teledyne Isco) with linear gradient from 0 to 20% phase B in 20 min (phase A, 0.02% trifluoroacetic acid in H_2_O; phase B, acetonitrile) and flow rate of 35 ml/min. PUM-containing fractions were pooled, concentrated, and lyophilized twice to yield 196 mg of a white solid highly soluble in water, DMSO, and methanol.

### Structure elucidation of PUM

Ion-trap ESI-MS (Bruker Esquire 3000 Plus showed a protonated molecular ion at *m/z* 487 [M+H]^+^ and a bimolecular ion at 973 [2M+H]^+^ Extended Data Fig. 1). Ion-trap ESI––MS/MS (Bruker Esquire 3000 Plus) showed major peaks at *m/z* 334, 353, 371, 389, 452, and 479 [M+H]^+^. HR-MS (Thermo Fisher Exactive) showed an exact mass of 487.18865, consistent with the molecular formula C_17_H_26_N_8_O_9_.

Reversed-phase HPLC (Shimadzu Series 10 with SPD-M10A VP diode array detector; Waters Symmetry Shield RP8 5μm, 250 x 4.6 mm, column; phase A = 2 mM heptafluorobutyric acid in water; phase B = 2 mM heptafluorobutyric acid in acetonitrile; gradient = 0% B at 0 min, 10% B at 20 min, 95% B at 30 min; flow rate = 1 ml/min) showed a single peak with a retention time of 12 minutes. The UV-absorbance spectrum showed maxima at 200 nm and 262 nm, consistent with the presence of a pyrimidine moiety^33^ (Extended Data Fig. 1).

The ^1^H NMR spectrum (600 MHz Bruker spectrometer in DMSO-d6 at 25°C) revealed one olefinic (H6), five amide (H6', H3, H5, Hα, and Hε-Gln), four methylene (Hβ-Gln, Hγ-Gln, Hα-Gly, and H5'), and five methine (H1', H2', H3', H4', and Hα-Gln) signals (Extended Data Fig. 2a). The 2D ^1^H–^13^C-HSQC and -HMBC NMR spectra (600 MHz Bruker spectrometer in DMSO-d6 at 25°C) identified five carboxyl-amide groups (C2, C4, C=O Gln, C=O Gly, and Cδ-Gln), two olefinic carbons (C1 and C6), four methine carbons belonging to a sugar ring (C1', C2', C3', and C4'), one other methine carbon (Cα-Gln), and four methylene groups (Cα-Gly, Cβ-Gln, Cγ-Gln, and C5') (Extended Data Fig. 2b-c).

The COSY NMR spectrum (600 MHz Bruker spectrometer in DMSO-d6 at 25°C) identified correlations between the five sugar protons: H1' (δH 4.35), H2' (δH 3.92), H3' (δH 3.65), H4' (δH 3.65) and H5' (δH-A 3.24; H-B 3.17). The chemical shift of C5' and ribose ^15^N-HSQC correlations between H6' and H5' indicated the presence of 6'-amino-ribose. Nitrogen signals N2 and N3 in the ^15^N HSQC spectrum, HMBC correlations of H1' to C1, C2 and C6, and HMBC correlations of H6 to C1', C1, C2, and C6, indicated the presence of uracil C-linked to C1' of 6'-amino-ribose. ^13^C-NMR spectra and COSY indicated the presence of a glutaminyl moiety, and HMBC correlations of H6' and the methine at δ_H_ 4.72 (Hα-Gln) to the carbonyl at δ_c_ 169.5 indicated linkage of the glutaminyl moiety to N6' of 6'-amino-ribose. Nε at δ 108.9 was assigned by ^15^N HSQC. The absence of a glutaminyl Nα signal in the ^15^N HSQC spectrum suggested the possible presence of an hydroxamic acid, and this was confirmed by reduction of PUM with aqueous TiCl_3_^34^ to yield desoxy-PUM (**1** in Extended Data Fig. 7a), exhibiting ion-trap ESI-MS mass of 471 [M+H]^+^, ion-trap ESI-MS/MS major peaks at *m/z* 355 and 372 [M+H]^+^, and a new nitrogen peak at δ117.7 in the ^15^N HSQC spectrum assignable as glutaminyl Nα, with corresponding NH at δH 8.34 coupling with δ_H_ 4.24 (Hα-Gln). The presence of two protons (Hα1 and Hβ2) coupling to a HN (δ_N_ 75) in COSY, the HMBC correlations of Hα1 and Hβ2 to a carbonyl at δ_c_ 157, and a NOESY NMR spectrum (600 MHz Bruker spectrometer in DMSO-d6 at 25°C) of desoxy-PUM indicated the presence of glycine C-linked to glutamine Nα. The chemical shift HN (δ_N_ 75) with corresponding δ_H_ 7.43 indicated the presence of formamidine C-linked to glycine Nα.

The stereochemistry of the glutamine residue was established to be (L) by total hydrolysis followed by chiral GC-MS (Hewlett-Packard HP5985B GC-MS; procedures as described^35^). The stereochemistry of the ribose sugar was inferred to be D by analogy to the natural product pseudouridine and was confirmed to be D by comparison of desoxy-PUM prepared by reduction of PUM with TiCl_3_ to desoxy-PUM prepared by total synthesis using a D-ribose precursor (Extended Data Fig. 7a-b).

### Effects of PUM on macromolecular synthesis

Cultures of *Staphylococcus simulans* strain M22 in 0.5x Mueller Hinton II broth (BD Biosciences) were incubated at 37°C with shaking until OD_600_ = 0.5; diluted in the same medium to OD_600_ = 0.1-0.2; supplemented with 6 kBq/ml [2-^14^C]-thymidine (Hartmann Analytic), 40 kBq/ml [5-^3^H]-uridine (Hartman Analytic), or 6 kBq/ml L-[^14^C(U)]-isoleucine (Hartman Analytic); and further incubated at 37°C with shaking. After 15 min, cultures were divided into two equal aliquots, PUM was added to one aliquot to a final concentration of 100-200 μM, and cultures were further incubated at 37°C with shaking. At time points 0, 10, 20, and 40 min following addition of PUM, 200 μl aliquots were removed, mixed with 2 ml ice-cold 10% TCA containing 1 M NaCl, and incubated 30-60 min on ice. The resulting TCA precipitates were collected by filtration on glass-microfiber filters (GF/C; Whatman), and filters were washed with 5 ml 2.5% TCA containing 1 M NaCl, transferred to scintillation vials, and dried. Filtersafe scintillation fluid was added (2 ml; Zinnser Analytic), and radioactivity was quantified by scintillation counting (Tri-Carb 3110TR Liquid Scintillation Analyzer; PerkinElmer).

### RNAP-inhibitory activity *in vitro*

For experiments in Fig. 1b and Extended Data Table 1 assessing promoter-dependent transcription by *E. coli* RNAP, reaction mixtures contained (25 µl): 0-20 µM PUM, 1 U *E. coli* RNAP σ^70^ holoenzyme (Epicentre), 10 nM DNA fragment carrying positions -112 to -1 of *E. coli recA* promoter^36^ followed by transcribed-region positions +1 to +363 of HeLaScribe Positive Control DNA (Promega; sequence at https://www.ncbi.nlm.nih.gov/pmc/articles/PMC4021065/bin/1471-2199-15-7-S1.docx; yields 363 nt transcript), 20 µM [α^32^P]GTP (0.3 Bq/fmol; PerkinElmer), 400 µM ATP, 400 µM CTP, and 6.25 μM, 50 μM, or 250 μM UTP, in 10 mM Tris-HCl (pH. 7.8), 2 mM HEPES-NaOH, 40 mM KCl, 3 mM MgCl_2_, 0.2 mM dithiothreitol, 0.09 mM EDTA, 0.2 mM phenylmethylsulfonyl fluoride, and 10% glycerol. Reaction components except RNAP were pre-equilibrated 10 min at 30°C. Reactions were initiated by addition of RNAP, were allowed to proceed 15 min at 30°C, and were terminated by addition of 175 µ1 HeLa Extract Stop Solution (Promega). Samples were phenol extracted and ethanol precipitated(procedures as described^37^), and pellets were resuspended in 10 μl 47.5% formamide, 10 mM EDTA, 0.025% bromophenol blue, and 0.01% xylene cyanol and heated 5 min at 95°C. Products were applied to 6% polyacrylamide (19:1 acrylamide:bisacrylamide; 7 M urea)^37^ slab gels, and gels were electrophoresed in TBE^37^ at 10 V/cm for 1.5 h, dried using a gel dryer (BioRad)^37^, and analyzed by storage-phosphor imaging (Typhoon; GE Healthcare).

For experiments in Fig. 1b and Extended Data Table 1 assessing promoter-dependent transcription by human RNAP I, human RNAP II, and human RNAP III, reaction mixtures contained (25 µl): 0-80 µM PUM, HeLa nuclear extract [3 μl HeLa nuclear extract prepared as described^38^, using ~4x10^7^ HeLa cells grown to 70-80% confluence in DMEM, high glucose, 2 mM L-glutamine medium containing 10% fetal bovine serum (Gibco) and 1% penicillin-streptomycin (Gibco) for assays of human RNAP I; or 6 U HeLaScribe Nuclear Extract (Promega) for assays of human RNAP II and human RNAP III], promoter DNA [4 nM *Eco*RI-linearized plasmid pHrP2x^39^ carrying human rDNA promoter for assays of human RNAP I (yields 379 nt transcript; 20 nM HeLaScribe Positive Control DNA (Promega) carrying cytomegalovirus immediate early promoter for assays of human RNAP II (yields 363 nt transcript with same sequence as *E.-coli-*RNAP-dependent transcript of preceding paragraph); or 2 nM plasmid pVAI^40^ carrying adenovirus VAI promoter for assays of human RNAP III (yields 160 nt transcript)], 20 µM [α^32^P]GTP (0.3 Bq/fmol: PerkinElmer), 400 µM ATP, 400 µM CTP, and 6.25, 50, or 250 µM UTP, in transcription buffer [12 mM HEPES-NaOH (pH 7.9), 75 mM KCl, 5 mM MgCl_2_, 10 mM creatine phosphate, 0.5 mM dithiothreitol, 0.1 mM EDTA, and 12% glycerol, for assays with human RNAP I; 10 mM Tris-HCl (pH 7.8), 2 mM HEPES-NaOH, 44 mM KCl, 3 mM MgCl_2_, 0.2 mM dithiothreitol, 0.09 mM EDTA, 0.2 mM phenylmethylsulfonyl fluoride, and 10% glycerol, for assays with human RNAP II and human RNAP III]. Procedures were as in the preceding paragraph.

For experiments in Fig. 1b assessing transcription by *B. subtilis* RNAP, SP6 RNAP, and T7 RNAP, , reaction mixtures contained (50 µl): 0-200 µM PUM, RNAP [0.2 U nM Rif-resistant *B. subtilis* [469 (513) Gln→Arg]β-RNAP σ^A^ holoenzyme (units defined as described^30^), 0.2 U SP6 RNAP (Promega), or 0.2 U T7 RNAP (Promega), DNA [0.2 nM plasmid pUC18 (Clontech; for assays with *B. subtilis* RNAP), or 0.2 nM plasmid pGEM-3Z (Promega; for assays with SP6 RNAP and T7 RNAP)], 500 µM ATP, 500 µM GTP, 500 µM CTP, and 6.25 μM [α^32^P]UTP (0.2 Bq/fmol; PerkinElmer; 6.25, 50, or 250 μM for Extended Data Table 1), in 40 mM Tris-HCl (pH 7.9), 6 mM MgCl_2_, 2 mM spermidine, 10 mM NaCl, 10 mM dithiothreitol, and 10 μg/ml bovine serum albumin. Reaction components except DNA were pre-equilibrated 15 min at 37°C. Reactions were initiated by addition of DNA, were allowed to proceed 15 min at 37°C, and were terminated by addition of 150 µl ice-cold 10% (w/v) TCA. After 30 min on ice, the resulting TCA precipitates were collected on glass-fiber filters (UniFilter GF/C; Perkin-Elmer) using a 96-well harvester (Packard). filters were washed once with water, and radioactivity was quantified using a TopCount scintillation counter (Packard).

For experiments in Fig. 1b assessing transcription by φ6 RNA-dependent RNAP, reaction mixtures contained (20 µl): 0-400 µM PUM, 0.5 U φ6 RNAP (Thermo Fisher), 2 µg poly(A) ssRNA (GE Healthcare), 1 µM poly(U-15) ssRNA primer (Sigma-Aldrich), 400 µM ATP, 400 µM GTP, 400 µM CTP, and 1.56 µM [α^32^P]UTP (0.02 Bq/fmol; Perkin-Elmer) in 50 mM Tris-acetate (pH 8.7), 50 mM ammonium acetate, and 1.5 mM MnCl_2_. Reaction components other than RNA template, RNA primer, and NTPs were pre-incubated 10 min at 30°C. Reactions were initiated by addition of RNA template, RNA primer, and NTPs, reactions were allowed to proceed 1 h min at 30°C, and reactions were terminated by spotting on DE81 filter discs (Whatman; pre-wetted with water) and incubating 1 min at 22°C. Filters were washed with 3x3 ml 0.5 M sodium phosphate dibasic, 2x3 ml water, and 3 ml ethanol using a filter manifold (Hoefer); filters were placed in scintillation vials containing 10 ml Scintiverse BD Cocktail (Thermo Fisher); and radioactivity was quantified by scintillation counting (LS6500; Beckman-Coulter).

For experiments in Extended Data Table 1 assessing promoter-independent transcription by *E. coli* RNAP and HeLa nuclear extract (human RNAP I/II/II), reaction mixtures contained (20 μl): 0-100 μM PUM, 100 nM *E. coli* RNAP core enzyme or 8 U HeLaScribe Nuclear Extract (Promega), 1 μg human placental DNA (Sigma-Aldrich; catalog number D7011), 400 μM ATP, 400 μM GTP, and 400 μM CTP, and 1.56, 25, or 400 μM [α^32^P]UTP (0.1-1 Bq/fmol; PerkinElmer), in 40 mM Tris-HCl (pH 8.0), 7 mM HEPES-NaOH, 70 mM ammonium sulfate, 30 mM KCl, 12 mM MgCl_2_, 5 mM dithiothreitol, 0.1 mM EDTA, and 12% glycerol. Procedures were performed as described^18^.

For experiments in Extended Data Figs. 4c and 7c, fluorescence-detected transcription assays were performed essentially as described^5^.

Half-maximal inhibitory concentrations (IC50s) were calculated by non-linear regression in SigmaPlot (Systat Software).

### Antibacterial activity *in vitro*

Antibacterial activities *in vitro* (Fig. 1c, rows 1-20; Extended Data Fig. 7d) were determined using broth-microdilution growth-curve assays^41^ (PUM degrades in phosphate-containing media with a half-life of ~12 h. Broth-microdilution endpoint assays^42^, which have a run time of 16-24 h^42^, which corresponds to 1.3 to 2 PUM half-lives, under-estimate absolute antibacterial activities of PUM. Broth-microdilution growth-curve assays^41^, which have shorter run times between assay start and assay signal, more accurately estimate absolute antibacterial activities of PUM.) Colonies of the indicated bacterial strains (5 to 10 per strain) were suspended in 3 ml phosphate-buffered saline^37^, suspensions were diluted to 1x10^5^ cfu/ml with growth medium [Todd Hewitt broth (BD Biosciences) for *Streptococcus* pyogenes and *Streptococcus pneumoniae*, aged Mueller Hinton II cation-adjusted broth (BD Biosciences; autoclaved and allowed to stand 2-12 months at room temperature before use) for *Staphylococcus aureus* and *Enterococcus faecium,* fresh Mueller Hinton II cation-adjusted broth (autoclaved and used immediately) for *Moraxella catarrhalis*, or fresh Mueller Hinton II cation-adjusted broth (BD Biosciences; autoclaved and used immediately) containing 0.4% *Haemophilus* Test Medium^43^ and 0.5% yeast extract for *Haemophilus influenzae*], 50 μl aliquots were dispensed into wells of a 96-well microplate containing 50 μl of the same medium or 50 μl of a 2-fold dilution series of PUM in the same medium (final concentrations = 0 and 0.25-256 μM), plates were incubated at 37°C with shaking, and optical densities at 600 nm were recorded at least hourly using a Synergy 2 (BioTek) or GENios Pro (Tecan) microplate reader. For each dilution series, growth curves were plotted, areas under growth curves were calculated, and IC50 was extracted as the lowest tested concentration of PUM that reduced area under the growth curve to 50% that in the absence of PUM (using only time points for rise phase of the growth curve in the absence of PUM).

Identical results were obtained in assays in the absence and presence of 30% human serum (Sigma-Aldrich; Fig. 1c, rows 1-4), indicating that PUM does not bind tightly to human serum proteins (unbound fraction ~100%).

Cytotoxicities for human macrovascular endothelial cells and human monocytes in culture (Fig. 1c, rows 21-22) were determined as described^44^.

### Antibacterial activity *in vivo*

Antibacterial activity *in vivo* was assessed in a mouse *S. pyogenes* peritonitis model (Fig. 1d; Extended Data Table 2). Female ICR mice (weight = 23-25 g; Harlan Laboratories Italy) were adapted to standardized environmental conditions (temperature = 23 ± 2°C; humidity = 55% ± 10%) for one week prior to infection. Infection was induced by intraperitoneal injection of 0.5 ml saline solution (supplemented with 1% peptone) containing 4 x 10^3^ cfu *S. pyogenes* C203 (an inoculum resulting in ≥95% mortality in untreated controls within 48 to 72 h after infection). Infected mice (eight mice per group; number determined by power calculations; assigned randomly to groups; unblinded) were treated with either: (i) 0.2 ml 5% dextrose or 0.2 ml of a 2.5-fold dilution series of PUM in 5% dextrose, administered intravenously 10 min after infection and again 6 h after infection (total PUM dose = 0 or 3.2-50 mg/kg), (ii) 0.25 ml 5% dextrose or 0.25 ml of a 2.5-fold dilution series of PUM in 5% dextrose, administered intravenously 10 min after infection (total PUM dose = 0 or 1.024-40 mg/kg), or (iii) or 0.25 ml 5% dextrose or 0.25 ml of a 2.5-fold dilution series of PUM in 5% dextrose, administered subcutaneously 10 min after infection (total PUM dose = 0 or 1.024-40 mg/kg). Survival was monitored for 7 days after infection. Experiments were performed in compliance with vertebrate animal ethical regulations and with Institutional Animal Care and Use Committee (IACUC) approval.

ED50s (doses yielding 50% survival at 7 days) and 95% confidence limits were calculated using the trimmed Spearman-Karber method as implemented in the US EPA LC50 Model System (http://sdi.odu.edu/model/lc50.php)^45^.

### Checkerboard interaction assays

Antibacterial activities of combinations of PUM and Rif were assessed in checkerboard interaction assays^46-47^ (Fig. 1e). Broth-macrodilution (assays procedures essentially as described^42^) were performed in checkerboard format, using *S. pyogenes* strain L49 or *S. pneumoniae* strain L44, and using Todd Hewitt broth (BD Biosciences) containing pairwise combinations of: (i) PUM at 1x, 0.5x, 0.25x, 0.125x, 0.063x, 0.031x, 0.016x, and 0.0078x MIC_PUM_ and (ii) Rif at 0.8x, 0.4x, 0.2x, 0.1x, 0.05x, 0.025x, and 0.0125x MIC_Rif_. Fractional inhibitory concentrations (FICs), FIC indices (FICIs), and minimum and maximum FICIs (FICI_min_ and FICI_max_) were calculated as described^47^. FICI_min_ ≤ 0.5 was deemed indicative of super-additivity (synergism), FICI_min_ > 0.5 and FICI_max_ ≤ 4.0 was deemed indicative of additivity, and FICI_max_ > 4.0 was deemed indicative of sub-additivity (antagonism)^46-47^.

### *S*pontaneous resistance rate assays

Spontaneous resistance rates were determined in Luria-Delbrück fluctuation assays^48^ (Fig. 2a). *S. pyogenes* strain ATCC 12344 (~1 x 10^9^ cfu/plate) was plated on Todd Hewitt agar [Todd Hewitt broth (BD Biosciences) supplemented with 1.5% Bacto agar (BD Biosciences)] containing 64 μg/ml or 128 μg/ml PUM (8x or 16x MIC under these conditions) or 1 μg/ml or 2 μg/ml Rif (8x or 16x MIC under these conditions), and numbers of colonies were counted after 24 h at 37°C (at least six independent determinations each). Resistance rates and 95% confidence intervals were calculated using the Ma-Sandri-Sarkar Maximum Likelihood Estimator^49-50^ as implemented on the Fluctuation Analysis Calculator (http://www.keshavsingh.org/protocols/FALCOR.html)^51^.

### Spontaneous PUM-resistant mutants, *S. pyogenes*

To isolate spontaneous PUM-resistant mutants of *S. pyogenes* (Fig. 2b), a single colony of *S. pyogenes* ATCC 12344 was inoculated into 5 ml Todd Hewitt broth (BD Biosciences) and incubated 3 h at 37°C with shaking in a 7% CO_2_/6% O_2_/4% H_2_/83% N_2_ atmosphere (atmosphere controlled using Anoxomat AN2CTS; Advanced Instruments), the culture was centrifuged, and the cell pellet (~1 x 10^9^ cells) was re-suspended in 50 μl Todd Hewitt broth and plated on Todd Hewitt agar (BD Biosciences) containing 16-256 μg/ml PUM (2-32x MIC under these conditions), and plates were incubated 120 h at 37°C in a 7% CO_2_/6% O_2_/4% H_2_/83% N_2_ atmosphere. PUM-resistant mutants were identified by the ability to form colonies on these media, were confirmed to be PUM-resistant by re-streaking on the same media, and were confirmed to be *S. pyogenes,* (as opposed to contaminants) using Taxo A differentiation discs (BD Biosciences) and Pyrase strips (Fluka).

Genomic DNA was isolated using the Wizard Genomic DNA Purification Kit [Promega; procedures as specified by the manufacturer, but with cells lysed using 1 mg/ml lysozyme (Sigma-Aldrich)] and was quantified by measurement of UV-absorbance (procedures as described^37^). The *rpoC* gene and the *rpoB* gene were PCR-amplified in reactions containing 0.2 μg genomic DNA, 0.4 μM forward and reverse oligodeoxyribonucleotide primers (5'-GGGCAAATGATAACTTAGTTGCGATTTGCTG-3' and 5'-CCTTTCTGCCTTTGATGACTTTACCAGTTC-3' for *rpoB;* 5'-GCTCAAGAAACTCAAGAAGTTTCTGAAACAACTGAC-3' and 5'-GTCAATGCTTTTTACTGCCAACAAACTCAGAC-3' for *rpoC*), 5 U Taq DNA polymerase (Genscript), and 800 μM dNTP mix (200 μM each dNTP; Agilent/Stratagene) (initial denaturation step of 3 min at 94°C; 30 cycles of 30 s at 94°C, 45 s at 53°C, and 4 min at 68°C; final extension step of 10 min at 68°C). PCR products containing the *rpoC* gene (3.6 kB) or the *rpoB* gene (3.6 kB) were isolated by electrophoresis on 0.8% agarose gels (procedures as described^37^), extracted from gel slices using a Gel/PCR DNA Fragments Extraction Kit (IBI Scientific; procedures as specified by the manufacturer), and sequenced (GENEWIZ; Sanger sequencing; seven sequencing primers per gene).

### Spontaneous PUM-resistant mutants, *E. coli*

To isolate spontaneous PUM-resistant mutants of *E. coli* (Extended Data Fig. 4a), *E. coli* uptake-proficient, efflux-deficient strain D21f2tolC^52^ was cultured to saturation in 10 ml LB broth^37^ at 37°C, cultures were centrifuged, cell pellets (~1 x 10^10^ cells) were re-suspended in 50 μl LB broth and plated on LB agar^37^ containing 800 μg/ml PUM (~1x MIC under these conditions), and plates were incubated 96-120 h at 37°C. PUM-resistant mutants were identified by the ability to form colonies on this medium.

Genomic DNA was isolated, and *rpoB* and *rpoC* genes were PCR-amplified and sequenced as described^18^.

### Resistance and cross-resistance levels

Resistance levels of *S. pyogenes* and *E. coli* spontaneous PUM-resistant mutants (Fig. 2b Extended Data Fig. 4b) were quantified in broth-microdilution assays. A single colony of a mutant strain or the isogenic wild-type parent strain was inoculated into 5 ml Todd Hewitt broth (BD Biosciences; for *S. pyogenes*) or LB broth^37^ (for *E. coli*) and incubated at 37°C with shaking in a 7% CO_2_/6% O_2_/4% H_2_/83% N_2_ atmosphere (atmosphere controlled using Anoxomat AN2CTS; Advanced Instruments); for *S. pyogenes*) or in air (for *E. coli*) until OD_600_ = 0.4-0.8. Diluted aliquots (~2 x 10^5^ cells in 50 μl same medium) were dispensed into wells of a 96-well microplate containing 50 μl of the same medium or 50 μl of a 2-fold dilution series of PUM in the same medium (final PUM concentration = 0 or 0.098-800 μg/ml), and were incubated 16 h at 37°C with shaking in a 7% CO_2_/6% O_2_/4% H_2_/83% N_2_ atmosphere (for *S. pyogenes*) or in air (for *E. coli*). MIC was defined as the lowest tested concentration of PUM that inhibited bacterial growth by ≥90%. MIC/MIC_wild-type_ was defined as the ratio of MIC for mutant to MIC for isogenic wild-type parent (*S. pyogenes* MIC_wild-type_ = 6.25 μg/ml under these conditions; *E. coli* MIC_wild-type_ = 400 μg/ml under these conditions).

Cross-resistance levels of *S. pyogenes* and *E. coli* spontaneous PUM-resistant mutants (Fig. 2e; Extended Data Fig. 4f) were determined as in the preceding paragraph, but using culture aliquots (~1x10^5^ cells) in 97 μl growth medium supplemented with 3 μl methanol or 3 μl of a 2-fold dilution series of Rif (Sigma-Aldrich; *S. pyogenes* MIC_wild-type_ = 0.098 μg/ml; *E. coli* MIC_wild-type_ = 0.20 μg/ml), lipiarmycin A3 (Lpm; BioAustralis; *S. pyogenes* MIC_wild-type_ = 6.25 μg/ml; *E. coli* MIC_wild-type_ = 1.56 μg/ml), myxopyronin B (Myx; prepared as described^53^; *S. pyogenes* MIC_wild-type_ = 6.25 μg/ml; *E. coli* MIC_wild-type_ = 0.20 μg/ml), streptolydigin (Stl; Sourcon-Padena; *S. pyogenes* MIC_wild-type_ = 3.13 μg/ml; *E. coli* MIC_wild-type_ = 3.13 μg/ml), CBR703 (CBR; Maybridge; *E. coli* MIC_wild-type_ = 6.25 μg/ml), or salinamide A (Sal; W. Fenical, Scripps Institution of Oceanography; *E. coli* MIC_wild-type_ = 0.049 μg/ml) in methanol (final concentrations = 0 and 0.006-50 μg/ml), or using culture aliquots (~2x10^5^) cells in 50 μl growth medium supplemented with 50 μl growth medium or 50 μl of a 2-fold dilution series of GE23077 (GE; prepared as described^54^; *E. coli* MIC_wild-type_ = 500 μg/ml) in growth medium (final concentrations = 0 and 125-8000 μg/ml).

Cross-resistance levels of *S. pyogenes* Rif-resistant mutants to PUM (Fig. 1e) were determined were determined as described for cross-resistance levels of *S. pyogenes* spontaneous PUM-resistant mutants, but analyzing a collection of 13 *S. pyogenes* spontaneous Rif-resistant mutants [isolated and sequenced using the same procedures used for isolation and sequencing of *S. pyogenes* PUM-resistant mutants (Methods, Spontaneous PUM-resistant mutants, *S. pyogenes*), but using Todd Hewitt agar containing 1-16x MIC Rif (0.1-2 μg/ml under these conditions)] and the isogenic wild-type parent, and analyzing a 2-fold dilution series of PUM (final concentration = 0 or 1.56-100 μg/ml).

Cross-resistance levels of *E. coli* Rif-, Lpm-, Myx-, and Sal-resistant mutants to PUM (Extended Data Fig. 4g-i,l) were determined as described for resistance levels of *E. coli* spontaneous PUM-resistant mutants, but analyzing a collection of *E. coli* D21f2tolC derivatives^18^ comprising four chromosomal Rif-resistant mutants, five chromosomal Lpm-resistant mutants, five chromosomal Myx-resistant mutants, five chromosomal Sal-resistant mutants, and the isogenic wild-type parent, and analyzing a 2-fold dilution series of PUM (final concentration = 0 or 25-1600 μg/ml).

Cross-resistance levels of *E. coli* Stl-, CBR-, and GE-resistant mutants to PUM (Extended Data Fig. 4j-k,n) were determined analogously, analyzing a collection of *E. coli* D21f2tolC pRL706 and *E. coli* D21f2tolC pRL663 derivatives^5,14,16^ comprising five plasmid-based Stl-resistant mutants, five plasmid-based CBR-resistant mutants, six plasmid-based GE-resistant mutants, and plasmid-based wild-type isogenic parents. Single colonies were inoculated into 5 ml LB broth containing 200 μg/ml ampicillin (Sigma-Aldrich), incubated at 37°C with shaking until OD_600_ = 0.4-0.8, supplemented with IPTG (Gold Bio) to a final concentration of 1 mM, and further incubated 1 h at 37°C with shaking. Diluted aliquots (~2 x 10^5^ cells in 50 μl LB broth containing 200 μg/ml ampicillin and 1 mM IPTG) were dispensed into wells of a 96-well microplate containing 50 μl of the same medium or 50 μl of a 2-fold dilution series of PUM in the same medium (final concentration = 0 or 25-4000 μg/ml), and were incubated 16 h at 37°C with shaking.

Amino-acid substitutions that confer PUM-resistance in the context of *S. pyogenes* RNAP were re-constructed and re-analyzed in the context of *E. coli* RNAP using an *E. coli* plasmid-based resistance assay (Extended Data Fig. 4b). Site-directed mutagenesis (QuikChange Site-Directed Mutagenesis Kit; Agilent) was used to construct plasmid pRL706^28^ and pRL663^55^ derivatives encoding *E. coli* RNAP β-subunit and β'-subunit derivatives having amino-acid substitutions that confer PUM-resistance in *S. pyogenes* (sequences from Fig. 2b). The resulting plasmids were introduced by transformation into *E. coli* strain D21f2tolC^52^, and resistance levels of transformants were determined using the procedures of the preceding paragraph.

### Formation of RNAP-promoter open complex

Experiments (Extended Data Fig. 4a) were performed essentially as described^12^. Reaction mixtures contained (20 μl): test compound (0 or 500 μM PUM, 2 μM Rif, or 100 μM Lpm), 40 nM *E. coli* RNAP σ^70^ holoenzyme, 10 nM Cy5-labelled DNA fragment carrying positions -40 to +15 of *lacUV5* promoter (*lacUV5*-[-40;+15]-Cy5^12^), and 100 μg/ml heparin, in 50 mM Tris-HCl (pH 8.0), 100 mM KCl, 10 mM MgCl_2_, 1 mM dithiothreitol, 10 μg/ml bovine serum albumin, and 5% glycerol. Reaction components other than DNA and heparin were incubated 10 min at 37°C; DNA was added and reactions were incubated 15 min at 37°C; and heparin was added and reactions were incubated 2 min at 37°C. Products were applied to 5% TBE-polyacrylamide slab gels (Bio-Rad), electrophoresed in TBE^37^, and analyzed by fluorescence scanning (Typhoon 9400).

### Nucleotide addition in transcription initiation

Experiments were performed essentially as described^5^, using reaction mixtures that contained no inhibitor, 500 μM PUM, 2 μM Rif, or 100 μM Lpm, and using 5 μM [α^32^P]UTP (3 Bq/fmol; PerkinElmer) (Extended Data Fig. 4b).

### Nucleotide addition in transcription elongation

Experiments were performed essentially as described^5^, using reaction mixtures that contained no inhibitor, 500 μM PUM, 2 μM Rif, or 100 μM Lpm (Extended Data Fig. 4c).

### Nucleotide addition at elevated NTP concentrations

Experiments (Fig. 3a) were performed essentially as described above for assays of promoter-dependent transcription by *B. subtilis* RNAP (Methods, RNAP-inhibitory activity *in vitro*), using reaction mixtures (50μl) that contained 0 or 6 µM PUM, 0.4 U *E. coli* RNAP σ^70^ holoenzyme (Epicentre), 0.4 nM plasmid pUC19 Clontech), 80 mM HEPES-KOH (pH 7.5), 80 mM KCl, 4 mM MgCl_2_, 0.1 mM EDTA, 5 mM dithiothreitol, 100 μg/ml bovine serum albumin, and either (i) 100 µM ATP, 100 µM [α^32^P]CTP (0.2 Bq/fmol; PerkinElmer), 100 µM GTP, and 10-500 µM UTP; (ii) 10-500µM GTP, 100 µM ATP, 100 µM CTP, and 2 µM [α^32^P]UTP (0.2 Bq/fmol; PerkinElmer); (iii) 100 µM GTP, 10-500 µM A, 100 µM CTP, and 2 µM [α^32^P]UTP (0.2 Bq/fmol); or (iv) 100 µM GTP, 100 µM ATP, 10-500 µM CTP, and 2 µM [α^32^P]UTP (0.2 Bq/fmol). The reaction time was 30 min at 37°C.

Relative nucleotide incorporation was defined as the ratio of nucleotide incorporation in the presence of PUM to nucleotide incorporation in the absence of PUM.

### Nucleotide addition on standard and "U-less cassette" templates

Experiments (Fig. 3b) were performed essentially as described in the preceding section (Methods, Nucleotide addition at elevated NTP concentrations), using reaction mixtures (50 μl) that contained 0-20 µM PUM, 0.4 U *E. coli* RNAP σ^70^ holoenzyme (Epicentre), 2 µM [α^32^P]CTP (0.2 Bq/fmol; PerkinElmer), 100 µM ATP, 100 µM GTP, and 5 µM UTP, in 40 mM Tris-HCl (pH 7.5), 10 mM MgCl_2_, 5 mM dithiothreitol, 15 mM KCl, 0.01 % Triton X-100, and 100 μg/ml bovine serum albumin, and either (i) 50 nM DNA fragment carrying positions -112 to +8 of *E. coli recA* promoter^36^ followed by 5'-CAGGGACAAGTTAGTTCGTTCAGCGACACGCGGCAACAAG-3' (directs incorporation of U, G, A, and C) or (ii) 50 nM DNA fragment carrying positions -112 to +8 of *E. coli recA* promoter followed by 5'-CAGGGACAAGGAGACCAACGCAGCGACACGCGGCAACAAG-3' ("U-less cassette"; directs incorporation of G, A, and C).). The reaction time was 60 min at 37°C. Relative nucleotide incorporation was defined as the ratio of nucleotide incorporation in the presence of PUM to nucleotide incorporation in the absence of PUM.

### Template-sequence specificity of inhibition by PUM: single-nucleotide-addition reactions

Template-sequence specificity of inhibition by PUM was assessed in single-nucleotide-addition experiments (Fig. 3c) using *E. coli* RNAP transcription elongation complexes assembled on the following nucleic-acid scaffolds (black, DNA; red, RNA; asterisk, ^32^P):

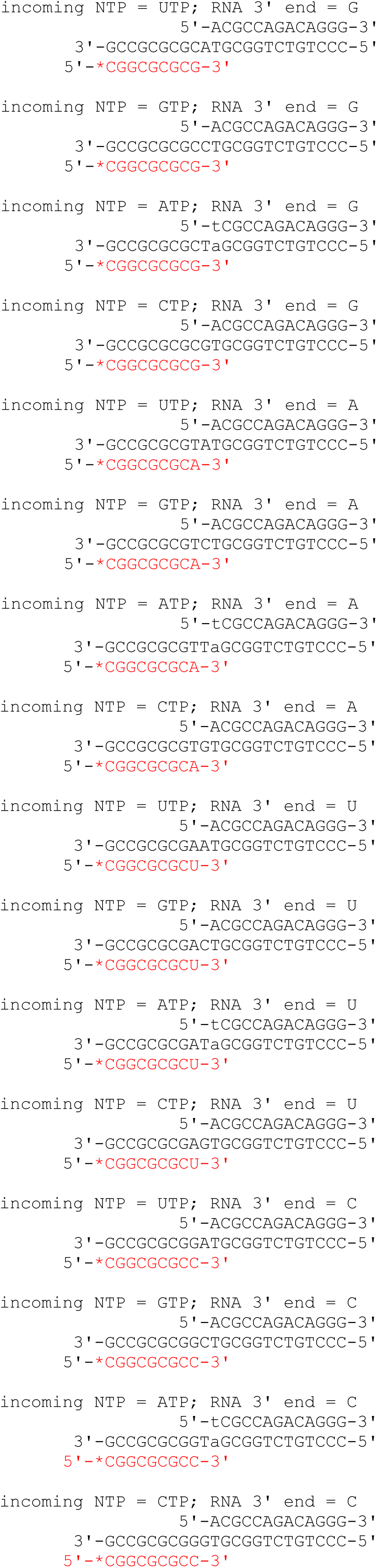

Nucleic-acid scaffolds for single-nucleotide-addition reactions were prepared as follows: 1 μM nontemplate-strand oligodeoxyribonucleotide [5'-ACGCCAGACAGGG-3' or 5'-TCGCCAGACAGGG-3'; IDT), 1 μM template-strand oligodeoxyribonucleotide [5'-GCCGCGCG-(C or T or A or G)-(A or C or G)-TGCGGTCTGTCCC-5' or 5'-GCCGCGCG-(C or T or A or G)-(T)-AGCGGTCTGTCCC-5'; IDT], and 1 μM ^32^P-5'-end-labeled oligoribonuceotide [5'-^32^P-CGGCGCGC-(U or C or A or G)-3'; 90 Bq/fmol; prepared using T4 polynucleotide kinase (New England Biolabs), [γ^32^P]ATP (100 Bq/fmol; PerkinElmer), and corresponding unlabelled oligoribonuceotide (IDT); procedures as described^37^], in 5 mM Tris-HCl (pH 7.7), 200 mM NaCl, and 10 mM MgCl_2_, were heated 5 min at 95°C, cooled to 4°C in 2°C steps with 1 min/step using a thermal cycler (Applied Biosystems), and stored at -20°C. Reaction mixtures for single-nucleotide-addition reactions contained (10 μl): 0 or 25 μM PUM, 40 nM *E. coli* RNAP core enzyme, 10 nM nucleic-acid scaffold, and 2.5 μM ATP, GTP, CTP, or UTP, in 50 mM Tris-HCl (pH 8.0), 100 mM KCl, 10 mM MgCl_2_, 1 mM dithiothreitol, 10 μg/ml BSA, and 5% glycerol. Reaction components except PUM and NTP were pre-incubated 10 min at 37°C, PUM was added and reaction mixtures were incubated 5 min at 37°C, and NTP was added and reaction mixtures were incubated 5 min at 37°C. Reactions were terminated by addition of 5 μl 80% formamide, 10 mM EDTA, 0.04% bromophenol blue, 0.04% xylene cyanol, and 0.08% amaranth red, and heating 2 min at 95˚C. Samples were applied to denaturing 15% polyacrylamide (19:1 acrylamide:bisacrylamide, 7 M urea)^37^ slab gels, electrophoresed in TBE^37^, and analyzed by storage-phosphor scanning (Typhoon 9400; GE Healthcare).

### Template-sequence specificity of inhibition by PUM: multiple-nucleotide-addition reactions

Template-sequence specificity of inhibition by PUM was assessed in multiple-nucleotide-addition experiments (Fig. 3d), performed by adding PUM to transcription elongation complexes halted at position +29 of a 100 bp transcription unit by omission of CTP, re-starting transcription elongation complexes and allowing transcription of positions +30 to +100 of the transcription unit by addition of CTP, and identifying positions at which PUM inhibits transcription.

Halted transcription elongation complexes were prepared as described^56^. Reaction mixtures (20 μl) contained: 40 nM *E. coli* RNAP σ^70^ holoenzyme, 10 nM DNA fragment N25-100-tR2^56^, 100 μg/ml heparin, 5 μM [γ^32^P]ATP (6 Bq/fmol; PerkinElmer), 5 μM UTP, and 5 μM GTP, in 50 mM Tris-HCl (pH 8.0), 100 mM KCl, 10 mM MgCl_2_, 1 mM dithiothreitol, 10 μg/ml bovine serum albumin, and 5% glycerol. Reaction components other than heparin and NTPs were pre-incubated 5 min at 30˚C; heparin was added and reaction mixtures were incubated 2 min at 30°C; NTPs were added and reaction mixtures were incubated 3 min at 30°C. Halted transcription elongation complexes were provided with PUM (1.25 μl 125 μM PUM, 1.25 μl 250 μM PUM, 1.25 μl 500 μM PUM, or 1.25 μl 1 mM PUM) or, to provide markers, chain-terminating 3'-O-methyl-NTPs (RiboMed; 1.25 μl 400 μM 3'-O-methyl-UTP, 1.25 μl 400 μM 3'-O-methyl-CTP, 1.25 μl 400 μM 3'-O-methyl-GTP, or 1.25 μl 400 μM 3'-O-methyl-ATP), were incubated 3 min at 30˚C, were re-started by addition of 0.625 μl 200 μM UTP, 1.25 μl 200 μM CTP, 0.625 μl 200 μM GTP, and 0.625 μl 200 μM ATP, and were further incubated 10 min at 30˚C. Reactions were terminated by addition of 12.5 μl 80% formamide, 10 mM EDTA, 0.04% bromophenol blue, 0.04% xylene cyanol, and 0.08% amaranth red, and heating 4 min at 95˚C. Samples were applied to 10% polyacrylamide (19:1 acrylamide:bisacrylamide; 7 M urea)^37^ slab gels, electrophoresed in TBE^37^, and analyzed by storage-phosphor scanning (Typhoon 9400).

### Structure determination: RPo-GpA-PUM

Crystals of *T. thermophilus* RPo-GpA were prepared as described^26^. Crystallization drops contained 1 μl 18 μM RPo (prepared from *T. thermophilus* RNAP σ^A^ holoenzyme and synthetic nucleic-acid scaffold as described^26^) and 1 mM GpA (RiboMed) in 20 mM Tris-HCl, pH 7.7, 100 mM NaCl, and 1% glycerol, and 1 μl reservoir buffer (RB; 100 mM Tris-HCl, pH 8.4, 200 mM KCl, 50 mM MgCl_2_, and 10% PEG4000), and were equilibrated against 400 μl RB in a vapor-diffusion hanging-drop tray (Hampton Research). Rod-like crystals appeared in 1 day, and grew to a final size of 0.1 mm x 0.1 mm x 0.3 mm in 5 days.

PUM was soaked into RPo-GpA crystals by adding 0.2 μl 100 mM PUM in RB to the crystallization drop and incubating 30 min at 22°C. RPo-GpA-PUM crystals were transferred to reservoir solutions containing 2 mM PUM in 17.5% (v/v) (2R,3R)-(-)-2,3-butanediol (Sigma-Aldrich) and were flash-cooled with liquid nitrogen.

Diffraction data for RPo-GpA-PUM were collected from cryo-cooled crystals at Cornell High Energy Synchrotron Source (CHESS) beamline F1. Data were integrated and scaled using HKL2000^57^. Structure factors were converted using the French-Wilson algorithm in Phenix^58^ and were subjected to anisotropy correction using the UCLA MBI Diffraction Anisotropy server (http://services.mbi.ucla.edu/anisoscale/)^59^.

The structure of RPo-GpA-PUM was solved by molecular replacement in Molrep^60^, using one RNAP molecule from the structure of *T. thermophilus* RPo (PDB 4G7H^26^) as the search model. Early-stage refinement included rigid-body refinement of each RNAP molecule, followed by rigid-body refinement of each subunit of each RNAP molecule. Cycles of iterative model building with Coot ^61^ and refinement with Phenix^62^ were performed. Atomic models of the DNA nontemplate strand, the DNA template strand, and GpA were built into mFo-DFc omit maps, and further refinement with Phenix was performed. The atomic model of PUM was built into the mFo-DFc omit map and was refined with Phenix. The final crystallographic model of RPo-GpA-PUM at 3.30 Å resolution, refined to Rwork and Rfree of 0.232 and 0.280, has been deposited in the PDB with accession code 5X21 (Fig. 4a; Extended Data Table 3).

### Structure determination: RPo-GpA-CMPcPP

Crystals of *T. thermophilus* RPo-GpA-CMPcPP were prepared by co-crystallization. Crystallization drops contained 1 μl 18 μM RPo (prepared from *T. thermophilus* RNAP σ^A^ holoenzyme and synthetic nucleic-acid scaffold as described^26^), 1 mM GpA (RiboMed), and 10 mM CMPcPP (Jena Bioscience) in 20 mM Tris-HCl, pH 7.7, 100 mM NaCl, and 1% glycerol, and 1 μl RB, and were equilibrated against 400 μl RB in a vapor-diffusion hanging-drop tray (Hampton Research). Rod-like crystals appeared in 1 day, and grew to a final size of 0.1 mm × 0.1 mm × 0.3 mm in 5 days. RPo-GpA-CMPcPP crystals were transferred to reservoir solutions containing 2 mM CMPcPP in 17.5% (v/v) (2R,3R)-(-)-2,3-butanediol (Sigma-Aldrich), and were flash-cooled with liquid nitrogen.

Diffraction data for RPo-GpA-CMPcPP were collected from cryo-cooled crystals at CHESS beamline F1. Data were integrated and scaled, structure factors were converted and subjected to anisotropy correction, and the structure was solved and refined using procedures analogous to those in the preceding section. The final crystallographic model of RPo-GpA-CMPcPP at 3.35 Å resolution, refined to Rwork and Rfree of 0.208 and 0.250, has been deposited in the PDB with accession code 5X22 (Fig. 4b; Extended Data Table 3).

### Semi-synthesis of PUM derivatives

Semi-syntheses of PUM derivatives from PUM corroborate the inferred structure of PUM, provide routes for preparation of novel PUM derivatives, and provide initial structure-activity relationships (Extended Data Fig. 7a,c-d). Reactions were conducted starting from 1 mg PUM, and products were identified by LC-MS (Agilent 1100 with flow split in 1:1 ratio between UV detector and ion-trap ESI-MS interface of Bruker Esquire 3000 Plus; Waters Atlantis 3 μm, 50 x 4.6 mm, column; phase A = 0.05% trifluoroacetic acid in water; phase B = acetonitrile; gradient = 5-95% B in 6 min; flow rate = 1 ml/min; run temperature = 40°C; PUM retention time = 1.4 min).

Reaction of PUM with TiCl_3_ in 1 M sodium acetate (pH 7.0) for 2 h at room temperature resulted in reduction of the N-hydroxy moiety, yielding desoxy-PUM (**1**; *m/z* = 471 [M+H]^+^). Reaction of PUM with PdCl_2_^63^ in 1:1 acetonitrile:water for 2 h at room temperature resulted in selective dehydration of the PUM glutamine sidechain amide -NH_2_, yielding nitrile analog **2** (*m/z* = 469 [M+H]^+^). Reaction of PUM with 0.1% TFA in water for 3 days at room temperature resulted in hydrolysis of the glutamine sidechain amide, yielding carboxy analog **3 (***m/z* = 488 [M+H]^+^). Reaction of **3** with benzylamine in DMF containing benzotriazol-1-yl-oxytripyrrolidinophosphonium hexafluorophosphate (PyBOP) for 30 min at room temperature yielded benzylamide analog **4** (*m/z* = 577 [M+H]^+^). Reaction of PUM with 2,3-butanedione^64^ in 10 mM ammonium acetate (pH 8.0) for 30 min at room temperature resulted in diolic intermediate **5** (*m/z* = 573 [M+H]^+^), which subsequently was trapped by treatment with phenylboronic acid for 2 h at room temperature, yielding phenyl-dioxaborolan analog **6** (*m/z* = 659 [M+H]^+^).

### Total synthesis of desoxy-PUM

Total synthesis of desoxy-PUM provides a reference compound that corroborates the inferred stereochemistry of PUM [by comparison of desoxy-PUM prepared by total synthesis (**1** in Extended Data Fig. 7b) to desoxy-PUM prepared by semi-synthesis from PUM (**1** in Extended Data Fig. 7a] and provides an additional route to novel PUM derivatives. Desoxy-PUM was obtained in eight steps by convergent synthesis from commercially available β-D-pseudouridine and glycyl-L-glutamine, as follows (Extended data Fig. 7b):

<u>Acetonide protection (reaction a in Extended Data Fig. 7b).</u> To a solution of β-D-pseudouridine (Berry & Associates; 400 mg, 1.64 mmol) and 2,2-dimethoxypropane (Sigma-Aldrich; 12 ml) in dimethylformamide (8 ml), concentrated HCl (80 μL) was added, and the reaction mixture was stirred 5 h at room temperature. After neutralization with 2.5 M NaOH, solvent was removed under vacuum. ^1^H-NMR (400 MHz, D_2_O, δ-H): 1.35 (s, 3H, CH_3_), 1.56 (s, 3H, CH_3_), 3.67 (dd, 1H, J = 12.2, 5.65 Hz, H-5’), 3.75 (dd, 1H, J = 12.2, 3.75 Hz, H-5’), 4.11 (dd, 1H, H-4’), 4.75 (m, 2H), 4.86 (m, 1H), 7.62 (s, 1H, H-6).

<u>Mesylation (reaction b in Extended Data Fig. 7b).</u> To a solution of the crude product of the preceding step (419 mg, 1.47 mmol) in pyridine (Sigma-Aldrich; 4.7 ml), methanesulfonyl chloride (Sigma-Aldrich: 95 μL, 1.23 mmol) was added with stirring at 0°C. The reaction mixture was stirred at room temperature until completeness (16 h). Solvent was removed by rotary evaporation, and the raw material was purified by flash chromatography on Combiflash (Teledyne ISCO), yielding 475 mg of a white powder (95% yield). ^1^H-NMR (400 MHz, acetonitrile-d_3_, δ-H): 1.32 (s, 3H, CH_3_), 1.54 (s, 3H, CH_3_), 4.33 (dd, 1H, J = 11 Hz, H-5'), 4.46 (dd, 1H, J = 11 Hz, H-5'), 4.20 (m, 1H), 4.72 (dd, 1H), 4.80 (m, 2H), 7.55 (s, 1H, H-6), 10.23 (sb, 1H, NH), 10.45 (sb, 1H, NH).

<u>Azidation (reaction c in Extended Data Fig. 7b).</u> To a solution of the product of the preceding step (475 mg) in dimethylformamide (24 ml), sodium azide (Sigma-Aldrich: 476 mg) was added, the reaction mixture was stirred 4 h at 100°C, and solvent was removed by rotary evaporation. ^1^H-NMR (400 MHz, acetonitrile-d_3_, δ-H): 1.30 (s, 3H, CH_3_), 1.50 (s, 3H, CH_3_), 3.52 (d, 2H, J = 5.3 Hz, H-5'), 4.04 (m, 1H, H-3'), 4.69 (dd, 1H, H-4'), 4.75 (d, 1H, J = 3.3 Hz, H-1'), 4.87 (dd, 1H, J = 3.3 Hz, H-2'), 7.58 (s, 1H, H-6).

<u>Azide reduction (reaction d in Extended Data Fig. 7b).</u> To a solution of the crude product of the preceding step (193 mg) in tetrahydrofuran (8.8 ml) and water (1.8 ml), 1 M trimethylphosphine in tetrahydrofuran (Sigma-Aldrich; 0.74 ml) was added, the reaction mixture was stirred 2 h at room temperature, and solvent was removed by rotary evaporation. ^1^H-NMR (400 MHz, D_2_O, δ-H): 1.47 (s, 3H, CH_3_), 168 (s, 3H, CH_3_), 3.40 (dd, 1H, H-5'), 3.49 (dd, 1H, H-5'),4.38 (m, 1H, H-4'), 4.90 (dd, 1H, H-1'), 4.94 (d, 1H, H-3'), 5.05 (dd, 1H, H-2'), 7.76 (s, 1H, H-6).

<u>Fmoc protection (reaction e in Extended Data Fig. 7b).</u> To a solution of the crude product of the preceding step (22 mg, 0.11 mmol) in dioxane (150 μL) and water (250 μL) sodium carbonate (26.5 mg) was added, followed by Fmoc chloride (Sigma-Aldrich; 31 mg, 1.3 eq), and the reaction mixture stirred overnight at room temperature. After addition of water (5 ml), the reaction was extracted with ethyl acetate (3 x 5 ml), the combined organic extracts were extracted with saturated sodium bicarbonate (3 x 5 ml), the combined aqueous extracts were acidified to pH 1 with 1 M HCl and extracted with ethyl acetate (3 x 5 ml), and the combined organic extracts were treated with sodium sulfate and evaporated to dryness, providing Fmoc-glycl-L-glutamine in quantitative yield. ^1^H-NMR (400 MHz, D_2_O, δ-H): 1.98 (m, 1H, Asn-β), 2.18 (m, 1H, Asn-β), 2.33 (m, 2H, Asn-γ), 3.90 (m, 2H, Gly-α), 4.23 (m, 1H), 4.31 (m, 1H), 4.47 (dd, 1H, Asn-α), 7.31 (m, 2H, Ar), 7.38 (m, 2H, Ar), 7.69 (m, 2H, Ar), 7.81 (m, 2H, Ar).

<u>Coupling, Fmoc deprotection, and formamidinylation (reactions f-h in Extended Data Fig. 7b).</u> To a solution of the product of the preceding step (20 mg) and the product of the azide-reduction reaction (30 mg, 1.1 eq) in dry dimethylformamide (1.5 ml), N,N′-dicyclohexylcarbodiimide (Sigma-Aldrich; 18 mg, 1.2 eq) and 1-hydroxybenzotriazole (Sigma-Aldrich; 19.5 mg, 2 eq) were added, and the reaction mixture was stirred overnight at room temperature, and the solvent was evaporated under reduced pressure. To a solution of the crude coupled product (12 mg) in dimethylformamide (800 μl), piperidine (200 μl) was added, and the reaction mixture was stirred 10 min at 25°C, the solvent was evaporated under reduced pressure, and the residue was washed with methylene chloride (2 x 5 ml). To a solution of the crude Fmoc-deprotected product (22 mg) in methanol (300 μl), 3,5-dimethylpyrazole-1-carboxamidine (Sigma-Aldrich; 45 mg, 10 eq) was added, and the reaction mixture was stirred overnight at room temperature, followed by 6 h under reflux at 65°C to complete the reaction. The solvent was evaporated under reduced pressure, and the solid residue was washed with methylene chloride (2 x 10 ml). ^1^H-NMR (400 MHz, D_2_O/CD_3_OD, δ-H): 1.33 (s, 3H, CH_3_), 1.54 (s, 3H, CH_3_), 2.01 (m, 1H, Asn-β), 2.17 (m, 1H, Asn-β), 2.37 (m, 2H, Asn-γ), 3.37 (m, 1H, H-5'), 3.65 (m, 1H, H-5'), 4.04 (s, 2H, Gly-α), 4.03 (m, 1H), 4.11 (m, 1H), 4.42 (m, 1H), 4.63 (m, 1H), 7.53 (s, 1H, H-6).

<u>Acetonide deprotection (reaction i in Extended Data Fig. 7b).</u> A solution of the crude product of the preceding step (17 mg) in acetic acid:water (7:3; 2 ml) was stirred overnight at room temperature and then heated to 50°C for 10 h under argon. The solvent was evaporated under reduced pressure, and the solid residue was washed with methylene chloride (2 x 5 ml) and methanol (2 ml), yielding a white solid that, when analyzed by LC-MS [performed as described for LC-MS of PUM (Methods, Structure Elucidation of PUM); retention time = 14 min], 1D- and 2D-NMR, was indistinguishable from desoxy-PUM obtained by reduction of PUM with TiCl_3_ (**1** in Extended Data Fig. 7a). ^1^H-NMR (600 MHz, DMSO-d_6_ /D_2_O, δ-H): 1.75 (m, 1H, Asn-β), 1.90 (m, 1H, Asn-β), 2.10 (m, 2H, Asn-γ), 3.29 (m, 2H, H-5'), 3.72 (m, 2H), 3.87 (s broad, 2H, Gly-α), 3.96 (m, 1H, H-2'), 4.24 (m, 1H, Asn-α), 4.40 (d, 1H, J=5.3 Hz, H-1'), 6.73 (s broad, CONH_2_), 7.32 (s broad, CONH_2_), 7.40 (s, 1H), 8.11 (t broad, 1H, NH), 8.34 (d broad, 1H, NH-Asn). ^13^C-NMR (DMSO-d_6_, δ-H): 28.4, 31.9, 41.4, 44.0, 53.0, 72.3, 73.7, 79.9, 81.6, 110.4, 141.5, 152.2, 158.2, 164.2, 168.0, 171.3, 173.7.

### Data analysis

Data for RNAP-inhibitory activities, growth-inhibitory activities, resistance, and cross-resistance are means of at least two technical replicates. Data for mouse infection models, resistance-rate assays, and checkerboard interaction assays are means and 95% confidence intervals for eight biological replicates, at least six biological replicates, and at least five technical replicates, respectively.

### Data availability

Atomic coordinates and structure factors for crystal structures of RPo-GpA-PUM and RPo-GpA-CMPcPP have been deposited in the Protein Data Bank with accession numbers 5X21 and 5X22. 16S rRNA gene sequences of PUM producer strains ID38640 and ID38673 have been deposited in GenBank with accession numbers JQ929050 and JQ929051.

## Extended Data Information

**Extended Data Figure 1.**
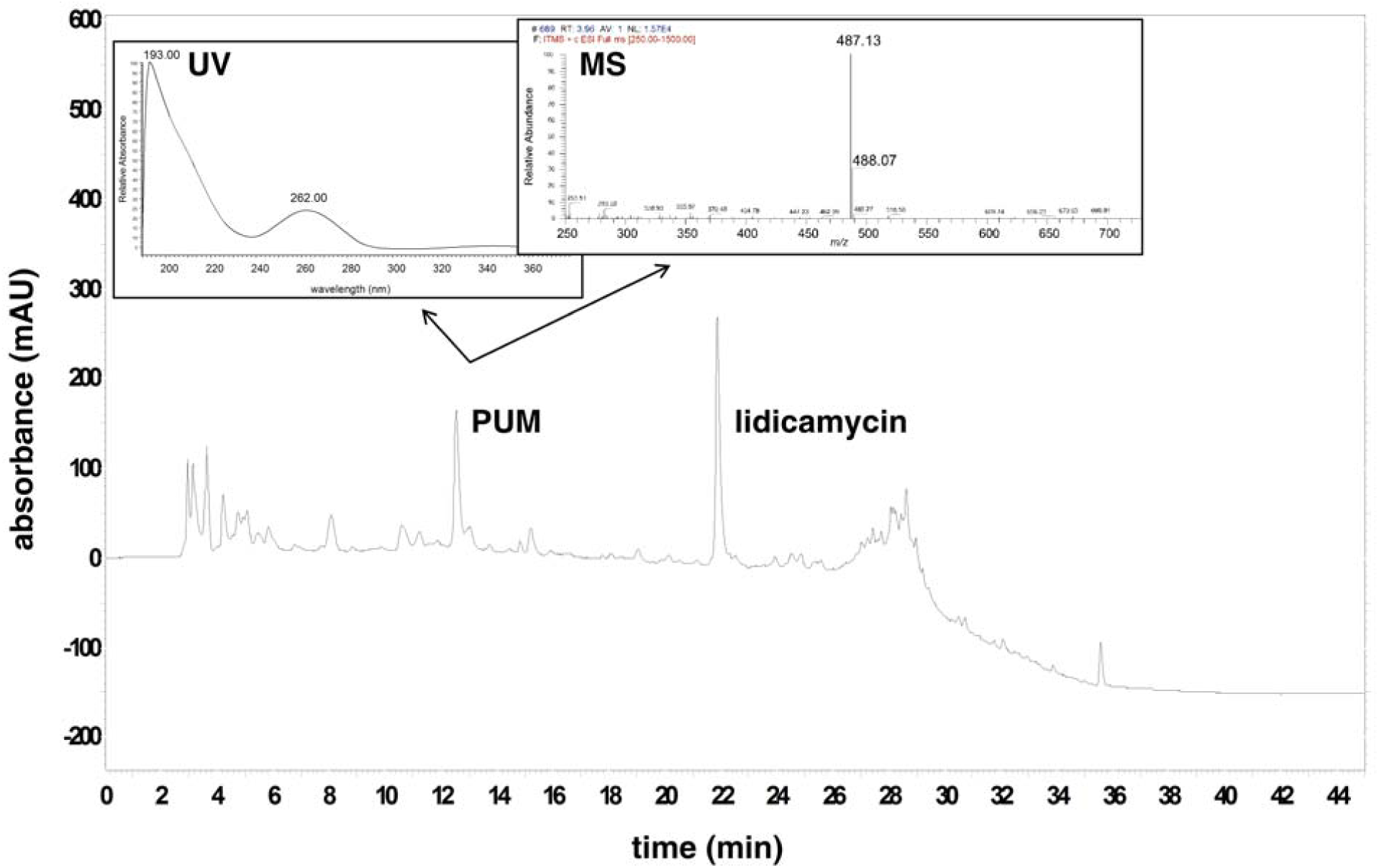
**Isolation of PUM.** Chromatographic profile of *Streptomyces* sp. ID38640 culture extract, showing peaks for PUM and lidicamycin, and UV-absorbance and mass spectra for PUM.

**Extended Data Figure 2.**
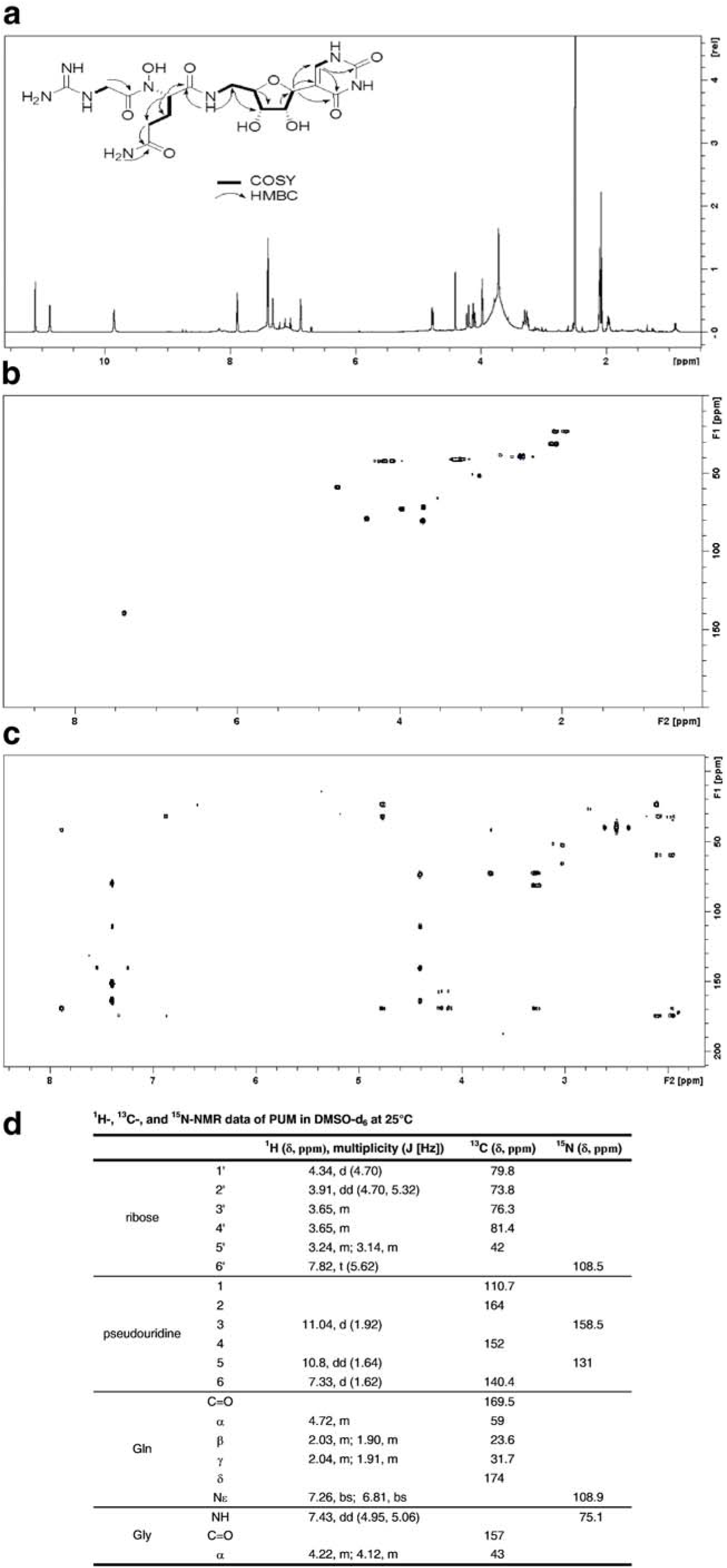
**Structure elucidation of PUM.a,** Structure and ^1^H-NMR spectrum of PUM in DMSO-d_6_ at 25°C at 400 MHz. **b,** 2D-HSQC spectrum of PUM in DMSO-d_6_ at 25°C at 400 MHz. **c,** 2D-HMBC spectrum of PUM in DMSO-d_6_ at 25°C at 400 MHz. **d,** Summary of ^1^H-,^13^C-, and ^15^N-NMR data for PUM in DMSO-d_6_.

**Extended Data Figure 3.**
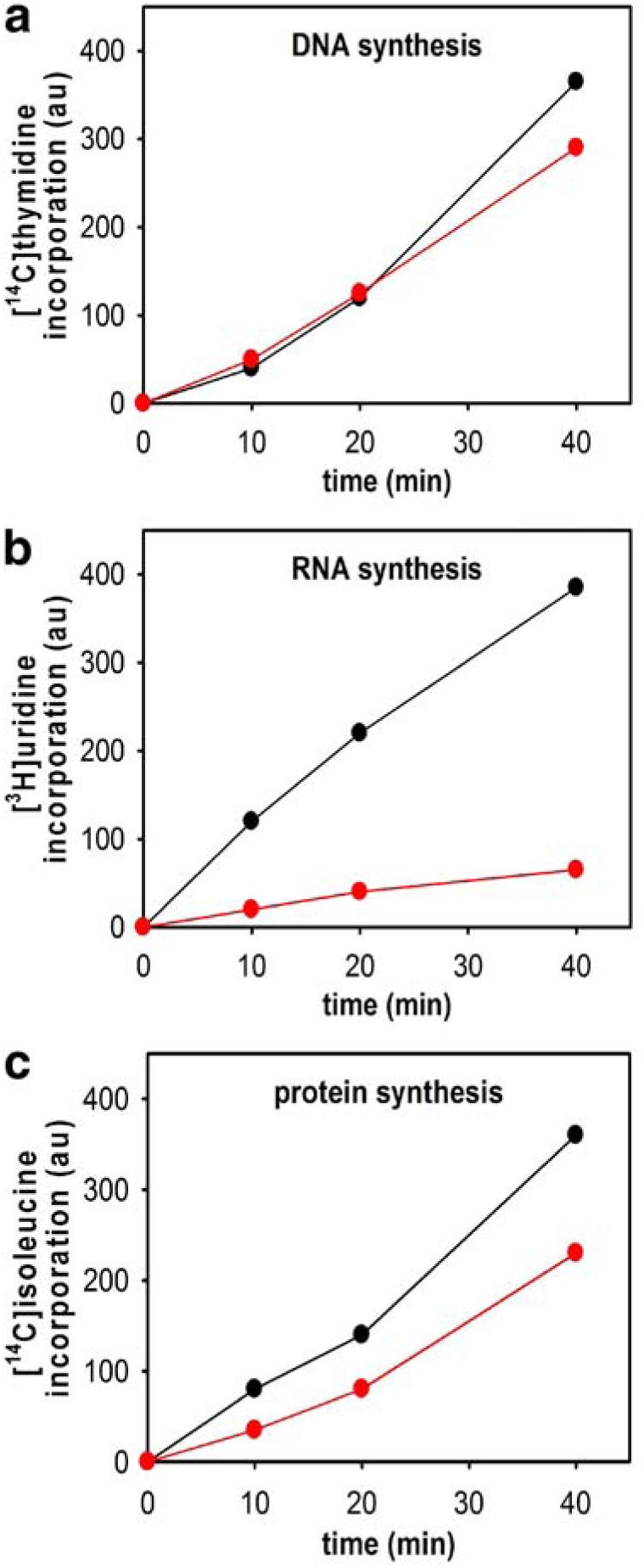
Effects of PUM on macromolecular synthesis in bacterial cells in culture: inhibition of RNAP-dependent RNA synthesis. Effects of PUM on DNA synthesis (a; [^14^C]-thymidine incorporation), RNA synthesis (b; [^3^H]-uridine incorporation), and protein synthesis (c; [^14^C]-isoleucine incorporation) in *Staphylococcus simulans* in culture. Results match characteristic pattern for inhibition of RNAP-dependent RNA-synthesis^18,65-70^: i.e., rapid and strong inhibition of RNA synthesis, slower and weaker inhibition of protein synthesis, and little or no inhibition of DNA synthesis.

**Extended Data Figure 4.**
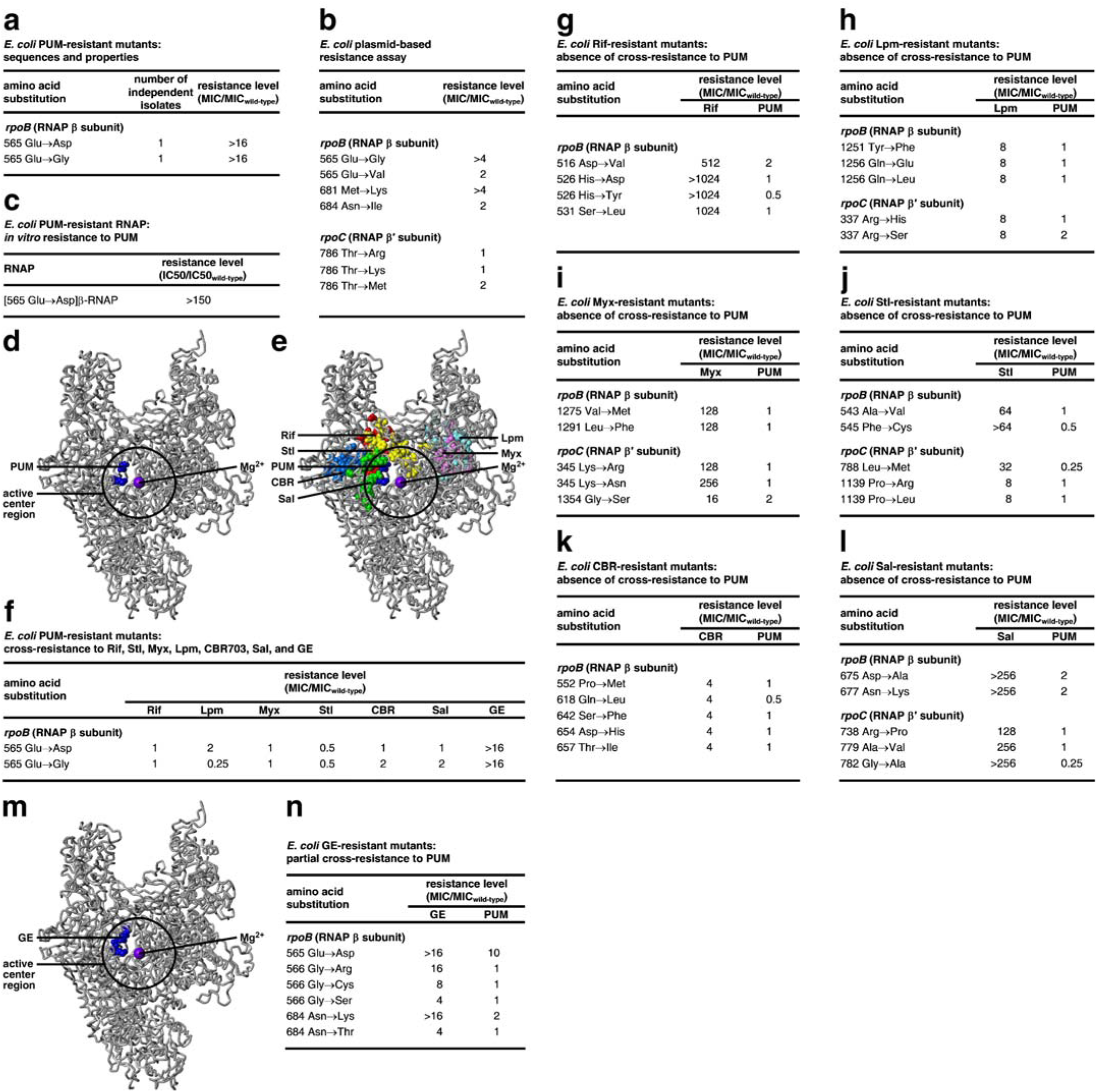
**Target of PUM: RNAP i+1 NTP binding site: results for Gram-negative bacterium *E. coli*. a,** *E. coli* spontaneous PUM-resistant mutants. **b,** Effects of *S. pyogenes* PUM-resistant mutants (sequences from Fig. 2b) when analyzed in *E. coli* plasmid-based resistance assay. Two substitutions confer moderate or higher (≥4x) resistance in *E. coli* plasmid-based resistance assay: β565 Glu→Gly and β681 Met→Lys. **c,** PUM-resistant phenotype of purified *E. coli* RNAP containing β565 Glu→Asp. **d,** Location of *E. coli* PUM target (sequences from **a**-**b**) in three-dimensional structure of bacterial RNAP (colors as in Fig. 2c). **e,** Absence of overlap between PUM target (blue) and Rif (red), Lpm (cyan), Myx (pink), Stl (yellow), CBR (light blue), and Sal (green) targets. **f,** Absence of cross-resistance of *E. coli* PUM-resistant mutants (sequences from **a**-**b**) to Rif, Lpm, Myx, Stl, CBR, and Sal. **g-l**, Absence of cross-resistance of *E. coli* Rif-, Lpm-, Myx-, Stl-, CBR-, and Sal-resistant mutants to PUM. **m,** Location of GE target (blue) in structure of bacterial RNAP. PUM target (**d**) shows partial overlap with GE target (**m**). **n,** Partial cross-resistance of *E. coli* GE-resistant mutants to PUM.

**Extended Data Figure 5.**
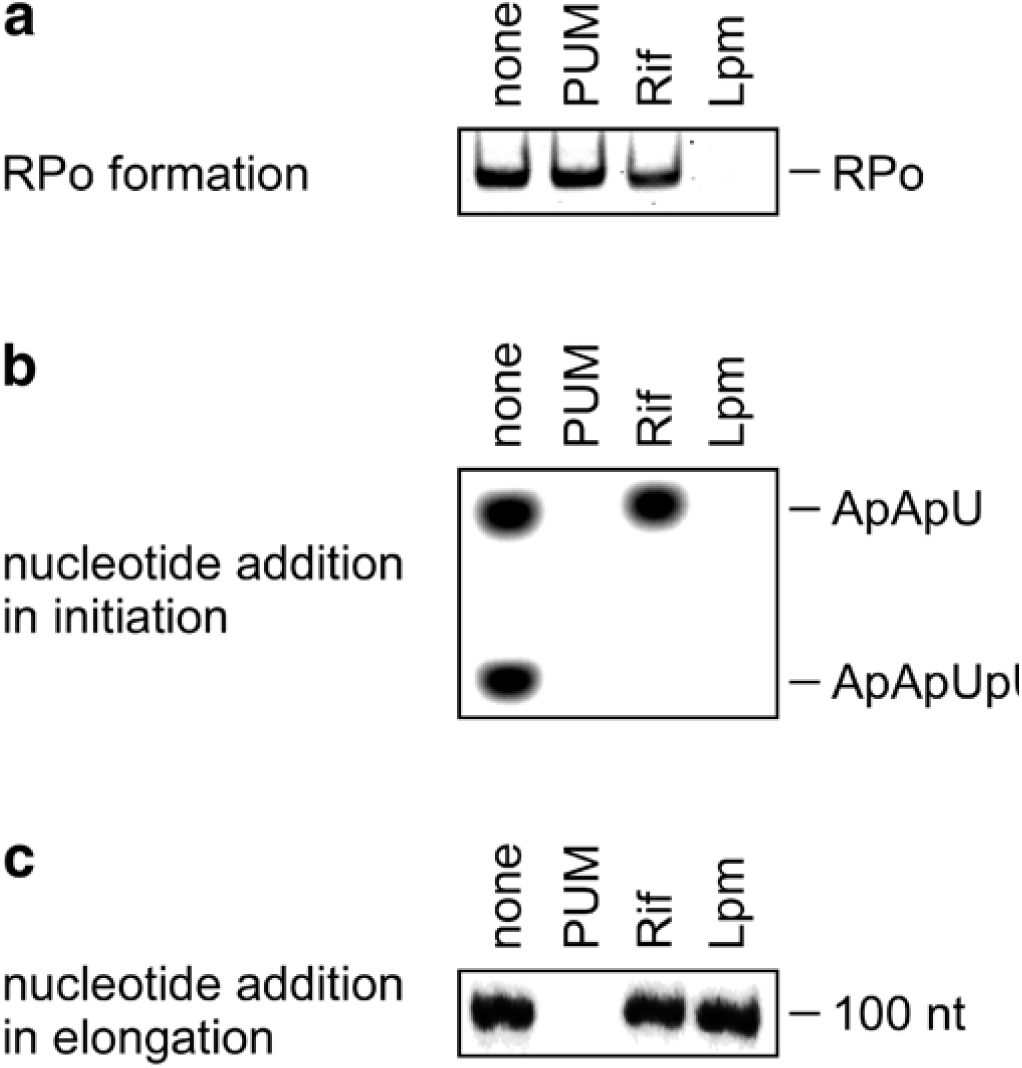
**Mechanism of PUM: inhibition of nucleotide addition a,** Absence of inhibition by PUM of formation of catalytically-competent RNAP-promoter open complex (RPo). **b,** Inhibition by PUM of nucleotide addition in transcription initiation. **c,** Inhibition by PUM of nucleotide addition in transcription elongation.

**Extended Data Figure 6.**
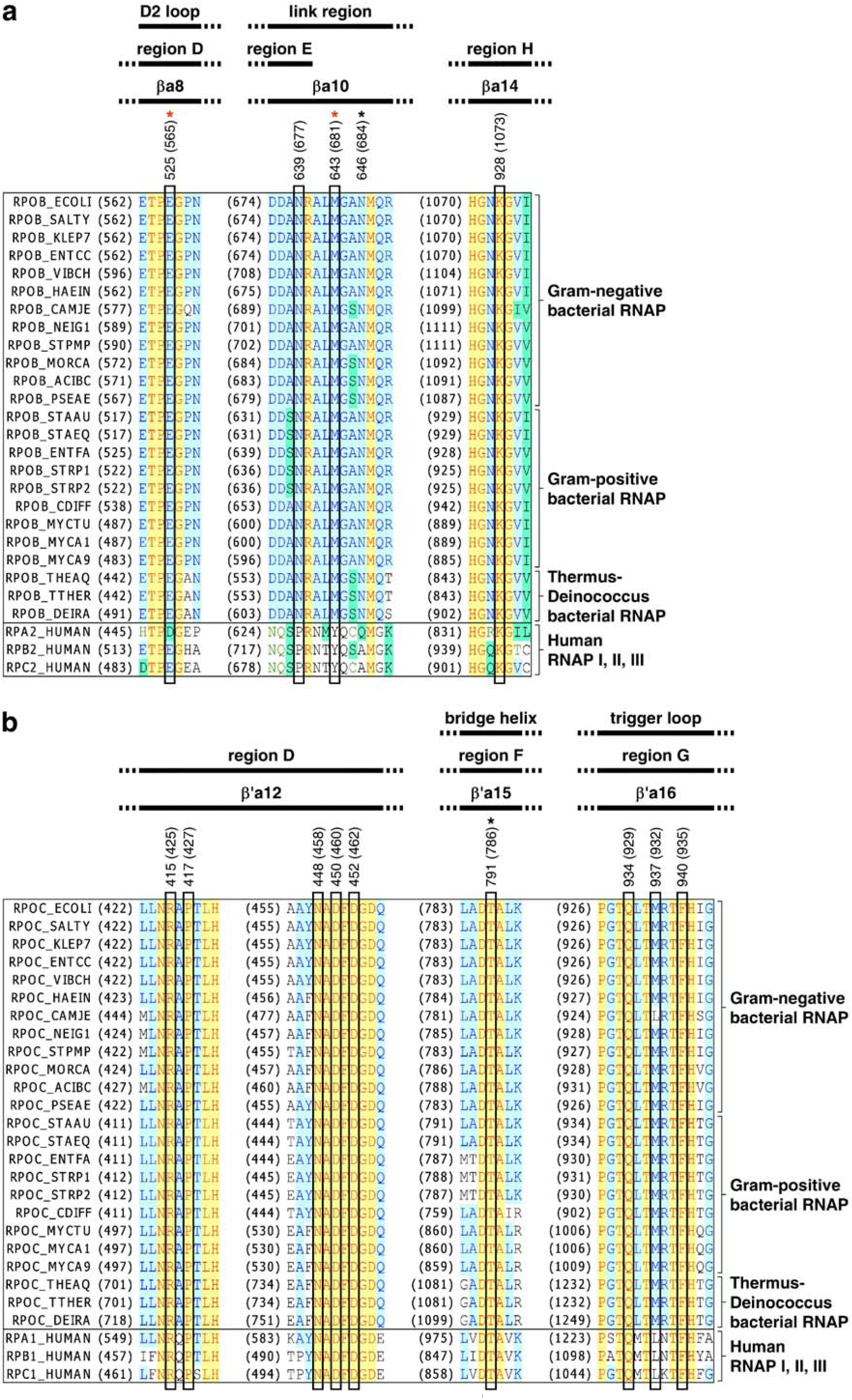
**Interactions between RNAP and PUM: sequence alignments.** Locations of residues that contact PUM in the sequences of RNAP β subunit (**a**) and RNAP β′ subunit (**b**). Sequence alignments for β and β′ subunits of bacterial RNAP (top 24 sequences in each panel) and corresponding subunits of human RNAP I, RNAP II, and RNAP III (bottom three sequences in each panel), showing locations of RNAP residues that contact PUM (black rectangles; numbered as in *S. pyogenes* and, in parentheses, as in *E. coli*; identities from Fig. 4a), locations of residues at which substitutions conferring PUM-resistance are obtained in both *S. pyogenes* and *E. coli* (red asterisks; identities from Fig. 2b and Extended Data Fig. 4a-b), locations of residues at which substitutions conferring PUM-resistance are obtained in *S. pyogenes* but not *E. coli* (black asterisks; identities from Fig. 2b and Extended Data Fig. 4a-b), locations of RNAP structural elements^71-72^ (top row of black bars), and RNAP conserved regions^73-75^ (next two rows of black bars). Species are as follows: *E. coli* (ECOLI), *Salmonella typhimurium* (SALTY), *Klebsiella pneumoniae* (KLEP7), *Enterococcus cloacae* (ENTCC), *Vibrio cholerae* (VIBCH), *Haemophilus influenzae* (HAEIN), *Campylobacter jejuni* (CAMJE), *Neisseria gonorrhoeae* (NEIG1), *Stenotrophomonas maltophilia* (STPMP), *Moraxella catarrhalis* (MORCA), *Acinetobacter baumannii* (ACIBC), *Pseudomonas aeruginosa* (PSEAE), *Staphylococcus aureus* (STAAU), *Staphylococcus epidermidis* (STAEQ), *Enterococcus faecalis* (ENTFA), *Streptococcus pyogenes* (STRP1), *Streptococcus pneumoniae* (STRP2), *Clostridium difficile* (CDIFF), *Mycobacterium tuberculosis* (MYCTU), *Mycobacterium avium* (MYCA1), *Mycobacterium abscessus* (MYCA9), *Thermus aquaticus* (THEAQ), *Thermus thermophilus* (THETH), *Deinococcus radiodurans* (DEIRA), and *Homo sapiens* (HUMAN).

**Extended Data Figure 7.**
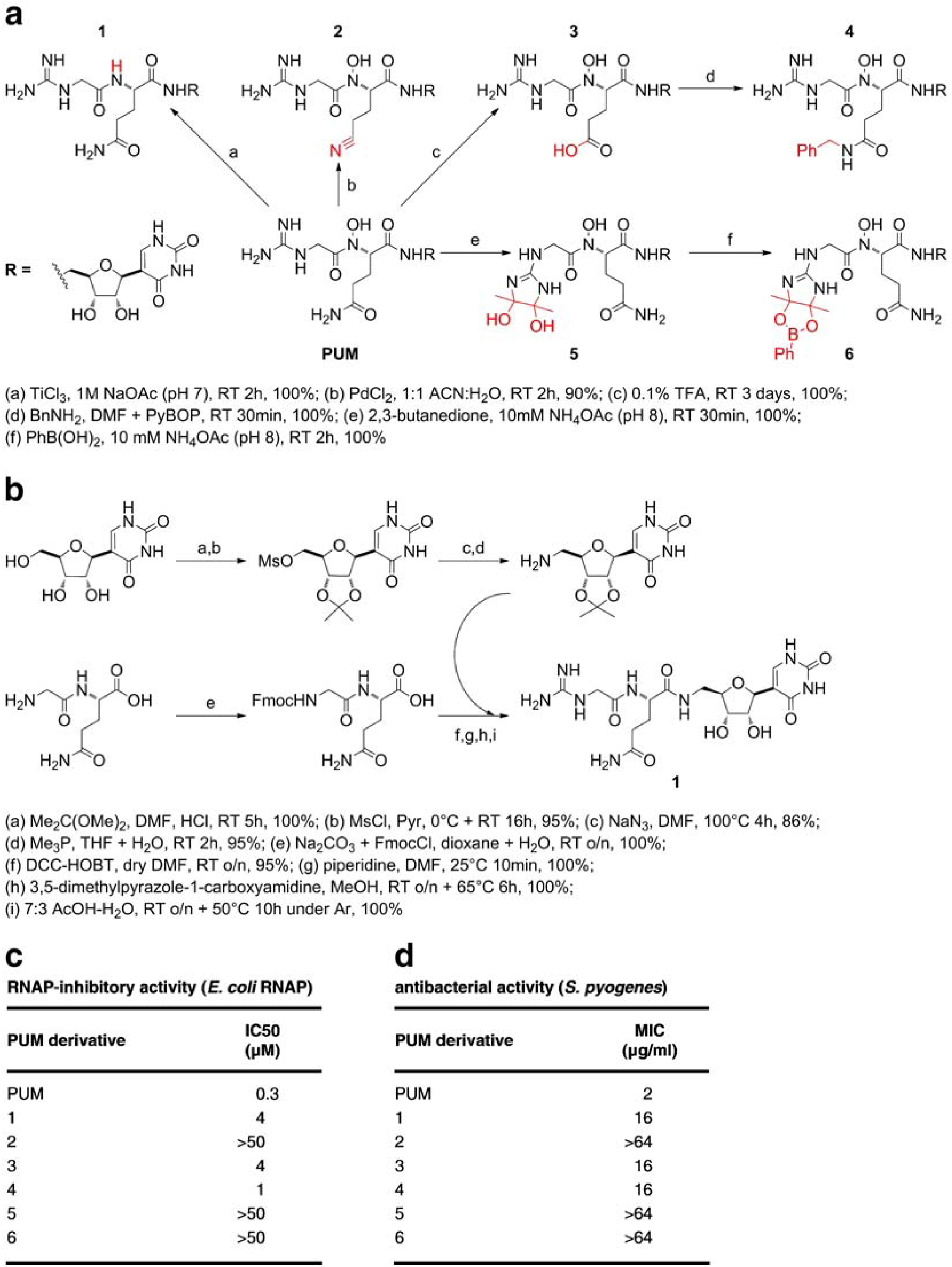
**Semi-synthesis, synthesis, and analysis of PUM derivatives. a,** Semi-synthesis of PUM derivatives lacking PUM N-hydroxy group (**1**), having alterations of PUM glutaminyl sidechain (**2**-**4**), or having alterations of PUM guanidinyl sidechain (**5**-**6**). **b,** Synthesis of PUM derivative lacking PUM N-hydroxy group (**1**). **c-d,** RNAP inhibitory activities and antibacterial activities of PUM derivatives.

**Extended Data Table 1.** Selectivity of RNAP-inhibitory activity

| enzyme | IC50 (μM) (± SEM) | selectivity ratio |
| --- | --- | --- |
| **promoter-dependent transcription** |  |  |
| ***E. coli* RNAP** |  |  |
| 6.25 μM UTP | 0.1 | [1] |
| 50 μM UTP | 2 | [1] |
| 250 μM UTP | 8 | [1] |
| **human RNAP I** |  |  |
| 6.25 μM UTP | >50 | >500 |
| 50 μM UTP | ND | ND |
| 250 μM UTP | ND | ND |
| **human RNAP II** |  |  |
| 6.25 μM UTP | 1 | 10 |
| 50 μM UTP | 7 | 4 |
| 250 μM UTP | >20 | >3 |
| **human RNAP III** |  |  |
| 6.25 μM UTP | 9 | 90 |
| 50 μM UTP | ND | ND |
| 250 μM UTP | ND | ND |
| **promoter-independent transcription** |  |  |
| ***E. coli* RNAP** |  |  |
| 1.56 μM UTP | 0.9 (± 0.1) | [1] |
| 25 μM UTP | 4 (± 0.7) | [1] |
| 400 μM UTP | 10 (± 2) | [1] |
| **HeLa nuclear extract (human RNAP I / II / III)** |  |  |
| 1.56 μM UTP | 4 (± 0.1) | 4 |
| 25 μM UTP | 15 (± 6) | 4 |
| 400 μM UTP | 51 (± 30) | 5 |

**Extended Data Table 2.** Antibacterial activity *in vivo* (mouse *S. pyogenes* peritonitis model; 7 day survival)

| intravenous (iv) administration,<br>10 min and 6 h post-infection |  | intravenous (iv) administration,<br>10 min post-infection |  | subcutaneous (sc) administration,<br>10 min post-infection |  |
| --- | --- | --- | --- | --- | --- |
| PUM total dose<br>(mg/kg) | survivors / total | PUM total dose<br>(mg/kg) | survivors / total | PUM total dose<br>(mg/kg) | survivors / total |
| 50 | 7 / 8 | 40 | 5 / 8 | 40 | 6 / 8 |
| 20 | 6 / 8 | 16 | 4 / 8 | 16 | 2 / 8 |
| 8 | 5 / 8 | 6.4 | 0 / 8 | 6.4 | 1 / 8 |
| 3.2 | 0 / 8 | 2.56 | 2 / 8 | 2.56 | 1 / 8 |
| 0 | 0 / 8 | 1.024 | 0 / 8 | 1.024 | 0 / 8 |
|  |  | 0 | 0 / 8 | 0 | 0 / 8 |

**Extended Data Table 3.** Data collection and refinement statistics.

|  | RPo-GpA-PUM<br>(5X21) | RPo-GpA-CMPcPP<br>(5X22) |
| --- | --- | --- |
| **Data collection** |  |  |
| Space group | C2 | P2<sub>1</sub> |
| Cell dimensions |  |  |
| *a*, *b*, *c* (Å) | 186.8, 103.1, 296.2 | 186.4, 104.3, 297.3 |
| $\alpha$, $\beta$, $\gamma$ (°) | 90.0, 98.7, 90.0 | 90.0, 98.4, 90.0 |
| Resolution (Å) | 40.00-3.30 (3.36-3.30)* | 50.00-3.35 (3.41-3.35)* |
| *R*<sub>sym</sub> or *R*<sub>merge</sub> | 0.138 (0.633) | 0.197 (>1.0) |
| *I*/σ(*I*) | 8.4 (1.7) | 7.7 (1.8) |
| Completeness (%) | 0.920 (0.874) | 0.956 (0.891) |
| Redundancy | 3.4 (3.4) | 5.3 (5.3) |
| **Refinement** |  |  |
| Resolution (Å) | 50.00-3.32 | 50.00-3.33 |
| No. reflections | 70085 | 147976 |
| *R*<sub>work</sub> / *R*<sub>free</sub> | 0.232/0.280 | 0.208/0.250 |
| Number of atoms |  |  |
| Protein/DNA/RNA | 28662 | 56787 |
| Ligand/ion | 42 | 76 |
| Water | 35 | 0 |
| *B* factors |  |  |
| Protein | 61.6 | 41.8 |
| Ligand/ion | 31.8 | 21.0 |
| Water | 34.3 | - |
| R.m.s deviations |  |  |
| Bond lengths (Å) | 0.002 | 0.007 |
| Bond angles (°) | 0.542 | 0.834 |

\*Values in parentheses are for highest-resolution shell.

## REFERENCES

1. Marston, H., Dixon, D., Knisely, J., Palmore, T. & Fauci, A. Antimicrobial resistance. JAMA 316, 1193–1204 (2016).

2. Brown, E. & Wright, G. Antibacterial drug discovery in the resistance era. Nature 529, 336–343 (2016).

3. Cihlar, T. & Ray, A. Nucleoside and nucleotide HIV reverse transcriptase inhibitors: 25 years after zidovudine. Antiviral Res. 85, 39–58 (2010).

4. Summers, B., Beavers, J. & Klibanov, O. Sofosbuvir [Solvadi], a novel nucleotide analogue inhibitor used for the treatment of hepatitis C virus. J. Pharm. Pharmacol. 66, 1653–1666 (2014).

5. Zhang, Y., Degen, D., Ho, M., Sineva, E., Ebright, K., Ebright, Y., Mekler, V., Vahedian-Movahed, H., Feng, Y., Yin, R., Tuske, S., Irschik, H., Jansen, R., Maffioli, S., Donadio, S., Arnold, E. & Ebright, R.H. GE23077 binds to the RNA polymerase 'i' and 'i+1' sites and prevents the binding of initiating nucleotides. eLife 3, e02450 (2014).

6. Landwehr, W., Wolf, C. & Wink, J. Actinobacteria and Myxobacteria: two of the most important bacterial resources for novel antibiotics. Curr. Top. Microbiol. Immunol. 398, 273–302 (2016).

7. Jin, D. J. & Gross, C. Mapping and sequencing of mutations in the Escherichia coli rpoB gene that lead to rifampicin resistance. J. Mol. Biol. 202, 45–58 (1988).

8. Garibyan, L., Huang, T., Kim, M., Wolff, E., Nguyen, A., Nguyen, T., Diep, A., Hu, K., Iverson, A., Yang, H. & Miller, J. Use of the *rpoB* gene to determine the specificity of base substitution mutations on the *Escherichia coli* chromosome, DNA Repair 2, 593–608 (2003).

9. Campbell, E., Korzheva, N., Mustaev, A., Murakami, K., Nair, S., Goldfarb, A. & Darst, S. Structural mechanism for rifampicin inhibition of bacterial RNA polymerase. Cell 104, 901–912 (2001).

10. Ebright, R. RNA exit channel–target and method for inhibition of bacterial RNA polymerase. WO/2005/001034 (2005).

11. Srivastava, A., Talaue, M., Liu, S., Degen, D., Ebright, R.Y., Sineva, E., Chakraborty, A., Druzhinin, S., Chatterjee, S., Mukhopadhyay, J., Ebright, Y., Zozula, A., Shen, J., Sengupta, S., Niedfeldt, R., Xin, C., Kaneko, T., Irschik, H., Jansen, R., Donadio, S., Connell, N. & Ebright, R.H. New target for inhibition of bacterial RNA polymerase: “switch region”. Curr. Opin. Microbiol. 14, 532–543 (2011).

12. Mukhopadhyay, J., Das, K., Ismail, S., Koppstein, D., Jang, M., Hudson, B., Sarafianos, S., Tuske, S., Patel, J., Jansen, R., Irschik, H., Arnold, E. & Ebright, R.H. The RNA polymerase "switch region" is a target for inhibitors. Cell 135, 295–307 (2008).

13. Belogurov, G., Vassylyeva, M., Sevostyanova, A., Appleman, J., Xiang, A., Lira, R., Webber, S., Klyuyev, S., Nudler, E., Artsimovitch, I. & Vassylyev, D. Transcription inactivation through local refolding of the RNA polymerase structure. Nature 45, 332–335. (2009).

14. Tuske, S., Sarafianos, S., Wang, X., Hudson, B., Sineva, E., Mukhopadhyay, J., Birktoft, J., Leroy, O., Ismail, S., Clark, A., Dharia, C., Napoli, A., Laptenko, O., Lee, J., Borukhov, S., Ebright, R.H. & Arnold, E. Inhibition of bacterial RNA polymerase by streptolydigin: stabilization of a straight-bridge-helix active-center conformation. Cell 122, 541–552 (2005).

15. Temiakov, D., Zenkin, N., Vassylyeva, M., Perederina, A., Tahirov, T., Kaihatsu, K., Savkina, M., Zorov, S., Nikiforov, V., Igarashi, N., Matsugaki, N., Wakatsuki, S., Severinov, K. & Vassylyev, D. Structural basis of transcription inhibition by antibiotic streptolydigin. Mol. Cell 19, 655–666 (2005).

16. Feng, Y., Degen, D., Wang, X., Gigliotti, M., Liu, S., Zhang, Y., Das, D., Michalchuk, T., Ebright, Y., Talaue, M., Connell, N. & Ebright, R.H. Structural basis of transcription inhibition by CBR hydroxamidines and CBR pyrazoles. Structure 23, 1470–1481 (2015).

17. Bae, B., Nayak, D., Ray, A., Mustaev, A., Landick, R. & Darst, S. CBR antimicrobials inhibit RNA polymerase via at least two bridge-helix cap-mediated effects on nucleotide addition. Proc. Natl. Acad. Sci. USA 112, E4178–E4187 (2015).

18. Degen, D., Feng, Y., Zhang, Y., Ebright, K., Ebright, Y., Gigliotti, M., Vahedian-Movahed, H., Mandal, S., Talaue, M., Connell, N., Arnold, E., Fenical, W. & Ebright, R.H. Transcription inhibition by the depsipeptide antibiotic salinamide A. eLife 3, e02451 (2014).

19. Sagitov, V., Nikiforov, V. & Goldfarb, A. Dominant lethal mutations near the 5' substrate binding site affect RNA polymerase propagation. J. Biol. Chem. 268, 2195–2202. (1993).

20. Svetlov, V., Vassylyev, D. & Artsimovitch, I. Discrimination against deoxyribonucleotide substrates by bacterial RNA polymerase. J. Biol. Chem. 279, 38087–38090 (2004).

21. Sosunov, V., Zorov, S., Sosunova, E., Nikolaev, A., Zakeyeva, I., Bass, I., Goldfarb, A., Nikiforov, V., Severinov, K. & Mustaev, A. The involvement of the aspartate triad of the active center in all catalytic activities of multisubunit RNA polymerase. Nucl. Acids Res. 33, 4202–4211 (2005).

22. Jovanovic, M., Burrows, P., Bose, D., Cámara, B., Wiesler, S., Weinzierl, R., Zhang, X., Wigneshweraraj, S. & Buck, M. An activity map of the *Escherichia coli* RNA polymerase bridge helix. J. Biol. Chem. 286, 14469–14479 (2011).

23. Yuzenkova, Y., Roghanian, M. & Zenkin, N. Multiple active centers of multi-subunit RNA polymerases. Transcription 3, 115–118 (2012).

## REFERENCES

1. Hudson, B., Quispe, J., Lara-González, S., Kim, Y., Berman, H., Arnold, E., Ebright, R. & Lawson, C. Three-dimensional EM structure of an intact activator-dependent transcription initiation complex. Proc. Natl. Acad. Sci. USA 106, 19830–19835 (2009).

2. Vrentas, C., Gaal, T., Ross, W., Ebright, R. & Gourse, R. Response of RNA polymerase to ppGpp: requirement for the ω subunit and relief of this requirement by DksA. Genes Dev. 19, 2378–2387 (2005).

3. Zhang, Y., Feng, Y., Chatterjee, S., Tuske, S., Ho, M., Arnold, E. & Ebright, R. H. Structural basis of transcription initiation. Science 338, 1076–1080 (2012).

4. Tang, H., Severinov, K., Goldfarb, A., Fenyo, D., Chait, B. & Ebright, R. Location, structure, and function of the target of a transcription activator protein. Genes Dev. 8, 3058–3067 (1994).

5. Severinov, K., Mooney, R., Darst, S. A. & Landick, R. Tethering of the large subunits of Escherichia coli RNA polymerase. J. Biol. Chem. 272, 24137–24140 (1997).

6. Niu, W., Kim, Y., Tau, G., Heyduk, T. & Ebright, R. Transcription activation at Class II CAP-dependent promoters: two interactions between CAP and RNA polymerase. Cell 87, 1123–1134 (1996).

7. Qi, Y. & Hulett, F. M. PhoP~P and RNA polymerase σA holoenzyme are sufficient for transcription of Pho regulon promoters in *Bacillus subtilis*: PhoP~P activator sites within the coding region stimulate transcription *in vitro.* Mol. Microbiol. 28 (1998).

8. Donadio, S., Monciardini, P. & Sosio, M. Approaches to discovering novel antibacterial and antifungal agents. Meths. Enzymol. 458, 3–28 (2009).

9. Mazza, P., Monciardini, P., Cavaletti, L., Sosio, M. & Donadio, S. Diversity of Actinoplanes and related genera isolated from an Italian soil. Microb. Ecol. 45, 362–372 (2003).

10. Ploeser, J. & Loring, H. The ultraviolet absorption spectra of the pyrimidine ribonucleosides and ribonucleotides. J. Biol. Chem. 178, 431–437 (1949).

11. Mattingly, P. & Miller, M. Titanium trichloride reduction of substituted N-hydroxy-2-azetidinones and other hydroxamic acids. J. Org. Chem. 45, 410–411 (1980).

12. Kettenring J., Colombo, L., Ferrari, P., Tavecchia, P., Nebuloni, M., Vékey, K., Gallo, G. & Selva, E. Antibiotic GE2270A: a novel inhibitor of bacterial protein synthesis: I. structure elucidation. J. Antibiot. 44, 702–715 (1991).

13. Sancar, A., Stachelejk, C., Konigsberg, W. & Rupp, Sequences of the recA gene and protein. Proc. Natl. Acad. Sci. USA 77, 2611–2615 (1980).

14. Sambrook, J. & Russell, D. Molecular Cloning: A Laboratory Manual (Cold Spring Harbor Laboratory, Cold Spring Harbor, NY, 2001).

15. Schreiber, E., Matthias, P., Müller, M. & Schaffner, W. Rapid detection of octamer binding proteins with 'mini-extracts', prepared from a small number of cells. Nucl. Acids Res. 17, 6419 (1989).

16. Pfleiderer, C., Smid, A., Bartsch, I. & Grummt I. An undecamer DNA sequence directs termination of human ribosomal gene transcription. Nucl. Acids Res. 18, 4727–4736 (1990).

17. Dean, N. & Berk, A. (1988) Ordering promoter binding of class III transcription factors TFIIIC1 and TFIIIC2. Mol. Cell. Biol. 8, 3017–3025.

18. Holowachuk, S., Bal'a, M. & Buddington, R. A kinetic microplate method for quantifying the antibacterial properties of biological fluids. J. Microbiol. Meth. 55, 441–446 (2003).

19. Clinical and Laboratory Standards Institute (CLSI/NCCLS). Methods for Dilution Antimicrobial Susceptibility Tests for Bacteria that Grow Aerobically; Approved Standard, Eighth Edition. CLIS Document M07-A8. Wayne, PA. (2009).

20. Barry, A., Pfaller, M. & Fuchs, P. Haemophilus test medium versus Mueller-Hinton broth with lysed horse blood for antimicrobial susceptibility testing of four bacterial species. Eur. J. Clin. Microbiol. Infect. Dis. 12, 548–553 (1993).

21. Mazzetti, C., Ornaghi, M., Gaspari, E., Parapini, S., Maffioli, S., Sosio, M. & Donadio, S. Halogenated spirotetronates from Actinoallomurus. J. Nat. Prod. 75 (2012).

22. Hamilton, M., Russo, R. & Thurston, R. Trimmed Spearman-Karber method for estimating median lethal concentrations in toxicity bioassays. Environ. Sci. Technol. 11, 714–719 (1977).

23. White, R., Manduru, M. & Bosso, J. Comparison of three different *in vitro* methods of detecting synergy: time-kill, checkerboard, and E test. Antimicrob. Agents Chemother. 40, 1914–1918 (1996).

24. Meletiadis, J., Pournaras, S., Roilides, E. & Walsh, T. Defining fractional inhibitory concentration index cutoffs for additive interactions based on self-drug additive combinations, Monte Carlo simulation analysis, and *in vitro-in vivo* correlation data for antifungal drug combinations against *Aspergillus fumigatus.* Antimicrob. Agents Chemother. 54, 602–609. (2010).

25. Srivastava, A., Degen, D., Ebright, Y. & Ebright, R. Frequency, spectrum, and nonzero fitness costs of resistance to myxopyronin in *Staphylococcus aureus.* Antimicrob. Agents Chemother. 56, 6250–6255 (2012).

26. Ma, W., Sandri, G. & Sarkar, S. Analysis of the Luria-Delbrück distribution using discrete convolution powers. J. Appl. Probab. 29, 255–267 (1992).

27. Sarkar, S., Ma, W. & Sandri, G. On fluctuation analysis: a new, simple and efficient method for computing the expected number of mutants. Genetica 85, 173–179 (1992).

28. Hall, B., Ma, C., Liang, P. & Singh, K. 2009. Fluctuation Analysis CalculatOR: a web tool for the determination of mutation rate using Luria-Delbrück fluctuation analysis. Bioinformatics 25, 1564–1565 (2009).

29. Fralick, J. & Burns-Keliher, L. Additive effect of tolC and rfa mutations on the hydrophobic barrier of the outer membrane of Escherichia coli K-12. J. Bacteriol. 176, 6404–6406 (1994).

30. Ebright, R. & Ebright, Y. Antibacterial agents: high-potency myxopyronin derivatives. WO/2012037508 (2013).

31. Ciciliato, I., Corti, E., Sarubbi, E., Stefanelli, S., Gastaldo, L., Montanini, N., Kurz, M., Losi, D., Marinelli, F. & Selva, E. Antibiotics GE23077, novel inhibitors of bacterial RNA polymerase I. taxonomy, isolation and characterization. J. Antibiot. 57, 210–217 (2004).

32. Wang, D., Meier, T., Chan, C., Feng, G., Lee, D. & Landick, R. Discontinuous movements of DNA and RNA in RNA polymerase accompany formation of a paused transcription complex. Cell 81, 341–350 (1995).

33. Revyakin, A., Liu, C., Ebright, R. & Strick, T. Abortive initiation and productive initiation by RNA polymerase involve DNA scrunching. Science 314, 1139–1143 (2006).

34. Otwinowski, Z. & Minor, W. Processing of X-ray diffraction data collected in oscillation mode. Meths. Enzymol. 276, 307–326 (1997).

35. French, S. & Wilson, K. On the treatment of negative intensity observations. Acta Cryst. D 34, 517–525 (1978).

36. Strong, M., Sawaya, M., Wang, S., Phillips, M., Cascio, D. & Eisenberg, D. Toward the structural genomics of complexes: crystal structure of a PE/PPE protein complex from *Mycobacterium tuberculosis.* Proc. Natl. Acad. Sci. USA 103 (2006).

37. Vagin, A. & Teplyakov, A. MOLREP: an automated program for molecular replacement. J. Appl. Cryst. 30 (1997).

38. Emsley, P., Lohkamp, B., Scott, W. & Cowtan, K. Features and development of Coot. Acta Cryst. D 66, 486–501 (2010).

39. Adams, P., Afonine, P., Bunkóczi, G., Chen, V., Davis, I., Echols, N., Head, J., Hung, L., Kapral, G., Grosse-Kunstleve, R., McCoy, A., Moriarty, N., Oeffner, R., Read, R., Richardson, D., Richardson, J., Terwilliger, T. & Zwart, P. PHENIX: a comprehensive Python-based system for macromolecular structure. Acta Cryst. D 66, 213–221 (2010).

40. Maffioli, S., Marzorati, E. & Marazzi. A. Mild and reversible dehydration of primary amides with PdCl2 in aqueous acetonitrile. Org. Lett. 7, 5237–5239 (2005).

41. Leitner, A. & Lindner, W. Probing of arginine residues in peptides and proteins using selective tagging and electrospray ionization mass spectrometry. J. Mass Spectrom. 38, 891–899 (2003).

42. Lancini, G. & Sartori, G. Rifamycins LXI: in vivo inhibition of RNA synthesis of rifamycins. Experentia 24, 1106–1106 (1968).

43. Lancini, G., Pallanza, R. & Silvestri, L. Relationships between bactericidal effect and inhibition of ribonucleic acid nucleotidyltransferase by rifampicin in *Escherichia* coli K-12. J. Bacteriol. 97, 761–768 (1969).

44. Sergio, S., Pirali, G., White, R. & Parenti F. Lipiarmycin, a new antibiotic from Actinoplanes III. Mechanism of action. J. Antibiot. 1975, 543–549 (1975).

45. Irschik, H., Gerth, K., Hofle, G., Kohl, W. & Reichenbach, H. The myxopyronins, new inhibitors of bacterial RNA synthesis from *Myxococcus fulvus* (Myxobacterales). J. Antibiot. 36, 1651–1658 (1983).

46. Irschik, H., Jansen, R., Hofle, G., Gerth, K. & Reichenbach, H. The corallopyronins, new inhibitors of bacterial RNA synthesis from Myxobacteria. J. Antibiot. 38, 145–152 (1985).

47. Irschik, H., Augustiniak, H., Gerth, K., Hofle, G. & Reichenbach, H. The ripostatins, novel inhibitors of eubacterial RNA polymerase isolated from Myxobacteria. J. Antibiot. 48, 787–792 (1995).

48. Weinzierl, R. The nucleotide addition cycle of RNA polymerase is controlled by two molecular hinges in the Bridge Helix domain. BMC Biol. 8, 134 (2010).

49. Hein, P. & Landick, R. The bridge helix coordinates movements of modules in RNA polymerase. BMC Biol. 8 (2010).

50. Sweetser, D., Nonet, M. & Young, R. Prokaryotic and eukaryotic RNA polymerases have homologous core subunits. Proc. Natl. Acad. Sci. USA 84, 1192–1196 (1987).

51. Lane, W. & Darst, S. Molecular evolution of multisubunit RNA polymerases: sequence analysis. J. Mol. Biol. 395, 671–685 (2010).

52. Jokerst, R., Weeks, J., Zehring, W., & Greenleaf, A. Analysis of the gene encoding the largest subunit of RNA polymerase II in Drosophila. Mol. Gen. Genet. 215, 266–275 (1989).

